# Discovery of decreased ferroptosis in male colorectal cancer patients with KRAS mutations

**DOI:** 10.1101/2023.02.28.530478

**Authors:** Hong Yan, Ronan Talty, Abhishek Jain, Yuping Cai, Jie Zheng, Xinyi Shen, Engjel Muca, Philip B. Paty, Marcus W. Bosenberg, Sajid A. Khan, Caroline H. Johnson

## Abstract

Aberrant tumor metabolism is a hallmark of cancer in which metabolic rewiring can support tumor growth under nutrient deficient conditions. KRAS mutations occur in 35-45% of all colorectal cancer (CRC) cases and are difficult to treat. The relationship between mutant KRAS and aberrant metabolism in CRCs has not been fully explored and could be a target for intervention. We previously acquired non-targeted metabolomics data from 161 tumor tissues and 39 normal colon tissues from stage I-III chemotherapy naïve CRC patients. In this study, we revealed that tumors from male patients with KRAS mutations only, had several altered pathways that suppress ferroptosis, including glutathione biosynthesis, transsulfuration activity, and methionine metabolism. To validate this phenotype, MC38 CRC cells (KRAS^G13R^) were treated with a ferroptosis inducer; RAS-selected lethal (RSL3). RSL3 altered metabolic pathways in the opposite direction to that seen in KRAS mutant tumors from male patients confirming a suppressed ferroptosis metabolic phenotype in these patients. We further validated gene expression data from an additional CRC patient cohort (Gene Expression Omnibus (GEO), and similarly observed differences in ferroptosis-related genes by sex and KRAS status. Further examination of the relationship between these genes and overall survival (OS) in the GEO cohort showed that KRAS mutant tumors are associated with poorer 5-year OS compared to KRAS wild type tumors, and only in male patients. Additionally, high compared to low expression of *GPX4, FTH1, FTL*, which suppressed ferroptosis, were associated with poorer 5-year OS only in KRAS mutant tumors from male CRC patients. Low compared to high expression of *ACSL4* was associated with poorer OS for this group. Our results show that KRAS mutant tumors from male CRC patients have suppressed ferroptosis, and gene expression changes that suppress ferroptosis associate with adverse outcomes for these patients, revealing a novel potential avenue for therapeutic approaches.

## Introduction

Colorectal cancer (CRC) is the third most diagnosed cancer worldwide. The KRAS oncogene, mutated in approximately 40% of CRC patients(1), is both a prognostic and predictive biomarker for which treatment is tailored. The incidence of tumors with KRAS mutations is higher on the right-side of the colon and occurs more frequently in female patients than males(2). Emerging evidence suggests that KRAS mutations induce metabolic reprogramming in CRC cells(3,4). This finding presents an opportunity to target tumor-specific metabolic vulnerabilities. Recent studies have revealed that CRC metabolism is dependent on the sex of the patient and cancer stage(5,6). However, no studies have reported on the association between the KRAS genotype and CRC metabolism in patients or examined sex differences in metabolic phenotypes related to KRAS.

Cancer cells undergo metabolic rewiring to fulfill the energy and biomass requirements of growing cells. Amino acids are metabolites and nutrients that are essential to the survival of all cell types, however they are also involved in reprogramming cancer metabolism. Cysteine, in particular, contributes to cellular fitness and survival by aiding growth in the tumor microenvironment and upon drug exposure(7). It is also an important intermediate in the transsulfuration pathway (TSP), a critical pathway that regulates ferroptosis; an iron-dependent, oxidative-stress-induced form of cell death that is tightly interwoven with cell metabolism(8). Depletion of cysteine triggers ferroptosis, however, to date, there is a lack of understanding regarding cysteine dependency and the role of other metabolites in KRAS mutant CRC. It is not known how KRAS-mediated alterations on metabolism are linked to ferroptosis and how these metabolic changes modulate signaling and CRC progression.

In this study, we show clear differences in tumor metabolism between KRAS mutant and KRAS wild type CRCs. We also show that ferroptosis related metabolites were altered in KRAS mutant tumors but only in those from male patients with CRC. We validated our findings using a gene expression dataset from another cohort, confirming these molecular differences in ferroptotic pathways are KRAS and sex-specific. In addition, we observed that genes which regulate ferroptosis associate with overall survival (OS), wherein high expression of these genes which suppress ferroptosis associate with poorer OS for male KRAS mutant patients only. This finding could contribute to the development of ferroptosis induction-based cancer therapy in a precision manner.

## Materials and methods

### Metabolomics Data collection and study design

We previously acquired non-targeted metabolomics data from 197 tumor tissues and 39 normal colon tissues(6). In our previous study 91 metabolites were identified as significantly different between normal colon and tumor tissues (6). Using recently updated clinical records, we further assigned these patients to KRAS mutant or KRAS wild type groups, in total 161 patients had KRAS genotype information and were assigned to either group. Detailed information for the clinical cohort is listed in **Supplementary Table 1**. KRAS mutations type for the patients are listed in **Supplementary Table 2**. The flowchart outlining of this study was summarized in **Figure 1**. A: Metabolomics data collection and analysis; including targeted analysis of metabolites involved in ferroptosis pathways; B: Identification of ferroptosis metabolic phenotype in RSL3 treated MC38 cell lines; C: Validation with transcriptomics data from a publicly available dataset.

**Figure 1.**
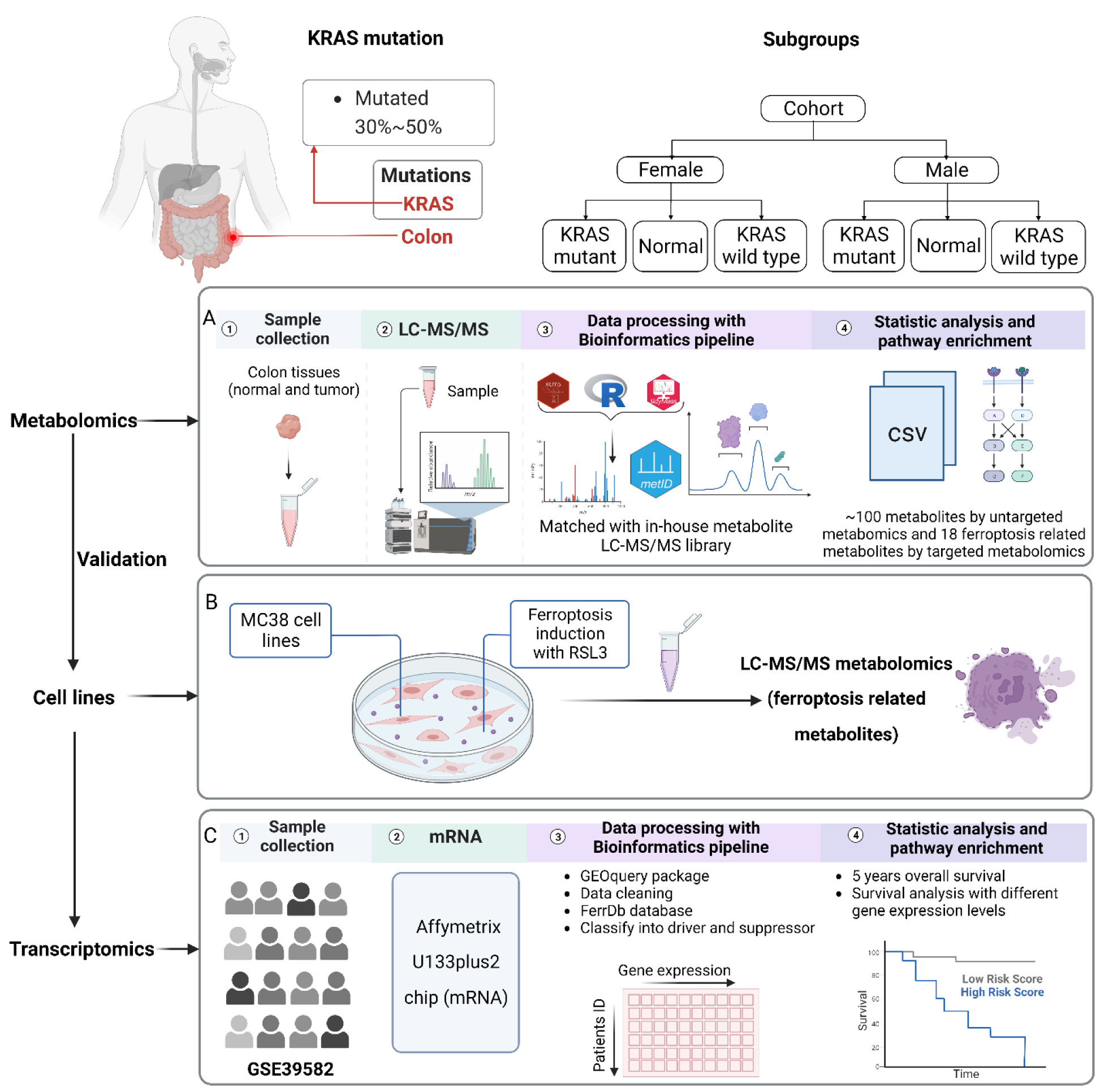
Flowchart outlining the study. A: Metabolomics data collection and analysis; including targeted analysis of metabolites involved in ferroptosis pathways; B: Identification of ferroptosis metabolic phenotype in RSL3 treated MC38 cell lines; C: Validation with transcriptomics data from a publicly available dataset.

**Table 1.** Differences in metabolite levels between KRAS mutant tumors, KRAS wild type tumors and normal tissues in male and female CRC patients. Wilcoxon Mann-Whitney U tests were performed and p-values were adjusted for false discovery rates (Benjamini-Hochberg). The p-values and fold changes that are significantly altered are in bold. Female patients: KRAS mutant type (n=35), KRAS wild type (n=43), normal (n=12). Male patients: KRAS mutant type (n=25), KRAS wild type (n=58), normal (n=27).

| Metabolites | Male adj.P value |  |  | Male Log2 fold change |  |  | Female adj.P value |  |  | Female Log2 fold change |  |  |
| --- | --- | --- | --- | --- | --- | --- | --- | --- | --- | --- | --- | --- |
|  | MT VS<br>WT | MT VS<br>normal | WT VS<br>normal | MT VS<br>WT | MT VS<br>normal | WT VS<br>normal | MT VS<br>WT | MT VS<br>normal | WT VS<br>normal | MT VS<br>WT | MT VS<br>normal | WT VS<br>normal |
| *xCT system* |  |  |  |  |  |  |  |  |  |  |  |  |
| Cystine | **5.00E-02** | 3.40E-01 | **5.30E-04** | **-0.44** | 1.19 | **1.63** | 5.50E-01 | 3.40E-01 | 1.20E-01 | -0.6 | 0.23 | 0.83 |
| Glutamine | 9.80E-01 | **1.00E-02** | **1.30E-04** | 0.1 | **0.53** | **0.43** | 9.50E-01 | **4.00E-02** | **6.90E-04** | 0.02 | **0.42** | **0.4** |
| Glutamate | 1.67E-01 | **1.15E-05** | **8.40E-12** | -0.12 | **1.12** | **1.24** | 5.35E-01 | **3.30E-03** | **2.96E-06** | -0.13 | **0.99** | **1.12** |
| GSH | 3.70E-01 | **5.00E-02** | 1.20E-01 | 0.1 | **2.34** | 2.25 | 7.30E-01 | **5.00E-02** | **1.00E-02** | 0.69 | **3.69** | **3** |
| GSSG | 4.10E-01 | 1.90E-01 | 2.40E-01 | 0.24 | 1.55 | 1.31 | 9.40E-01 | 2.10E-01 | 7.00E-02 | -0.1 | 1.25 | 1.35 |
| GSH/GSSG ratio | 8.60E-01 | **3.00E-02** | **1.00E-02** | 0.36 | **0.96** | **0.6** | 5.10E-01 | 6.00E-02 | 1.70E-01 | 0.36 | 0.96 | 0.6 |
| *TSP and BH4 metabolism* |  |  |  |  |  |  |  |  |  |  |  |  |
| SAH | 8.50E-01 | **2.60E-03** | **7.20E-05** | 0.01 | **0.92** | **0.91** | 6.00E-01 | **4.80E-03** | **7.00E-07** | -0.17 | **2.37** | **2.54** |
| SAM | 6.00E-02 | **1.00E-02** | **2.70E-03** | 0.17 | **0.16** | **-0.01** | 6.70E-01 | 1.10E-01 | **5.00E-02** | -0.11 | -0.28 | **-0.18** |
| SAM/SAH ratio | 7.20E-01 | **5.00E-02** | 1.40E-01 | 0.3 | **-0.67** | -0.98 | 1.20E-01 | **2.00E-02** | **4.00E-03** | 1.15 | **-0.62** | **-1.76** |
| Methionine | 2.20E-01 | **1.00E-02** | **1.60E-06** | -0.12 | **0.62** | **0.74** | 8.90E-01 | **2.00E-02** | **5.50E-04** | -0.04 | **0.12** | **0.16** |
| Serine | 4.00E-01 | **8.70E-04** | **1.00E-07** | -0.1 | **0.76** | **0.86** | 9.50E-01 | **1.00E-02** | **4.20E-05** | 0.01 | **0.6** | **0.59** |
| Dihydrobiopterin | **1.00E-02** | **2.00E-02** | **6.80E-07** | **-0.77** | **0.32** | **1.09** | 5.60E-01 | **1.00E-02** | **1.80E-06** | 0.62 | **2.21** | **1.59** |
| *LOX and COX pathways* |  |  |  |  |  |  |  |  |  |  |  |  |
| Adrenic acid | 9.70E-01 | 7.00E-02 | **2.00E-02** | 0 | 0.36 | **0.35** | 8.10E-01 | 4.10E-01 | 2.40E-01 | -0.07 | -0.07 | 0 |
| 5-HETE | 4.00E-01 | 1.00E-01 | **1.60E-03** | 0.08 | -0.26 | **-0.34** | 8.00E-02 | 2.60E-01 | **2.70E-03** | -0.81 | -0.36 | **0.45** |
| Prostaglandin E2 | 7.00E-01 | **5.00E-02** | 6.00E-02 | 0.45 | **0.9** | 0.46 | 6.50E-01 | 4.30E-01 | 2.90E-01 | -0.64 | 0.05 | 0.69 |
| Prostaglandin F2 $\beta$ | 1.50E-01 | **2.00E-02** | 1.00E-01 | 0.03 | **0.06** | 0.02 | 9.00E-02 | 7.00E-02 | 1.60E-01 | -0.49 | -2.14 | -1.65 |
| Arachidonic acid | 1.77E-01 | 1.42E-01 | 2.56E-01 | -0.14 | -0.10 | 0.03 | 3.26E-01 | 1.55E-01 | 1.93E-01 | -0.16 | -0.24 | -0.08 |
| *Mevalonate and Cholesterol metabolism* |  |  |  |  |  |  |  |  |  |  |  |  |
| GPP | 6.30E-01 | 2.00E-01 | **2.00E-02** | 0.15 | -0.02 | **-0.17** | 1.10E-01 | 1.10E-01 | 2.80E-01 | -0.13 | -0.32 | -0.2 |
| 7-Dehydrocholesterol | 8.20E-01 | **2.90E-03** | **8.50E-06** | -0.13 | **0.68** | **0.81** | 6.00E-02 | 8.00E-02 | **2.30E-04** | -0.43 | 0.37 | **0.81** |

WT: KRAS wild type; MT: KRAS mutant type; Adj.P value: p values adjusted for false discovery rates (FDR)
Abbreviations: GSH: Glutathione; GSSG, Glutathione disulfide; SAM, S-adenosyl-methionine; SAH, S-adenosyl-homocysteine; 5-HETE, 5-Hydroxyeicosatetraenoic acid; GPP, Geranyl pyrophosphate; TSP, Transsulfuration pathway; BH4, Tetrahydrobiopterin

#### Semi-targeted metabolomics of metabolites in ferroptosis pathways

Using one of the stored tumor or normal tissue homogenates, we extracted metabolites as described above. We also replicated the LC-MS conditions as aforementioned, performing MS/MS on both samples and commercial standards (Sigma Aldrich, St. Louis, MO and Cayman Chemical, Ann Arbor, MI). This step was carried out to validate the relative quantification and identity of the metabolites, which were housed in ferroptosis pathways.

### MC38 cell lines, conditions, and treatments

MC38 CRC cells were used in this study and obtained from NIH DTP, DCTD repository. Cells were cultured in DMEM/F12 (Thermo Fisher, #11320-033) including L-glutamine, and 2.438g/L sodium bicarbonate, and supplemented with 10 % fetal bovine serum (Thermo Fisher, #16140-071), 1x non-essential amino acids (Thermo Fisher #11140050), and 1x Penicillin and Streptomycin (Thermo Fisher, #15140-122). MC38 cells were seeded on 15 cm plates with five replicates one day prior to the experiment. On the following day, cells were treated with 10 μM RSL3 (1S,3R)-RSL3 (RSL3) (Cayman Chemical, Ann Arbor, MI) for 18 hours. All cells were cultured at 37 °C, 5% CO2 and kept at low passage (∼3-5 passages). Trypan Blue was used to assess cell count and viability after 18 hours of treatment, and normalization was performed in the reconstitution step for extract preparation based on cell count results. Collected cell lines were transferred to a −80 °C freezer until further usage in 2 mL Eppendorf tubes.

Prior to metabolomics, multiple experiments were conducted to confirm that RSL3 induces ferroptosis in MC38 cells. Cell viability was assessed via AquaBluer assay (Multitarget Pharmaceuticals, Colorado Springs, CO) after 24 hours of treatment with a range RSL3 doses (0 to 50 μM)(**Supplementary Figure 1A**). Cell viability was also assessed via AquaBluer assay after 24 hours treatment of 10 μM RSL3 treatment with and without 100 μM ferrostatin-1, a specific ferroptosis inhibitor (**Supplementary Figure 1B**). Finally, cellular lipid peroxidation was measured by treating cells with 10 μM RSL3 for 18 hours and staining with 10 μM Liperfluo (Dojindo, Rockville, MD) for 30 minutes with subsequent analysis by flow cytometry (**Supplementary Figure 1C). Supplementary Figure 1A** showed that 10 μM is around the LC50. 100 μM ferrostatin was chosen simply to validate that RSL3 is inducing ferroptotic cell death. Doses as low as 1 μM result in 100% cell death rescue (not shown).

### MC38 Cell metabolite extract preparation

A volume (500 μL) of ice-cold ACN: MeOH: H_2_O(2:2:1, v/v/v) was added to each sample as the extraction solvent. The samples were vortexed for 30 s and sonicated for 10 min. To precipitate proteins, the samples were incubated for 2 hours at −20 °C, followed by centrifugation at 13,000 rpm (15,000 g) and 4 °C for 15 min. The resulting supernatant was transferred to new 1.5 mL microcentrifuge tube and evaporated to dryness for 12 hours using a vacuum concentrator (Thermo Fisher Scientific, Waltham, MA, United States). The extraction was repeated and the dry extracts were then reconstituted in 100 μL of ACN:H2O (1:1, v/v), sonicated for 10 min, and centrifuged at 13,000 rpm (15,000 g) and 4 °C for 15 min to remove insoluble debris. The supernatant was transferred to UPLC autosampler vials (Waters cooperation, MA, United States). A pooled quality control (QC) sample was prepared by mixing 5 μL of extracted solution from each sample into a UPLC autosampler vial. All the vials were capped and stored at −80 °C prior to UPLC-MS analysis. MS analysis and the data preprocessing workflow are provided in Supplementary material.

### Gene expression data collection

The gene expression microarray dataset GSE39582 was retrieved from NCBIs gene expression omnibus (GEO) database (https://www.ncbi.nlm.nih.gov/geo/). The dataset contains mRNA expression profiles (Affymetrix U133Plus2) from 585 CRCs and 19 non-tumoral colorectal mucosal tissues. Within this dataset, tumor tissue samples were selected from stage I-III patients (n = 395) aged ≥55 years old; information regarding KRAS type, and sex of patient can be observed in **Supplementary Table 3**. Relevant to this study, this cohort contains gene expression information on *GPX4*, *FTH1*, *FTL* and *ACSL4* and clinical outcomes and KRAS status information for 174 females and 221 males with colon adenocarcinoma.

### Statistical analysis

Statistical analyses were performed in R (version 4.1.0). Nonparametric Kruskal–Wallis rank sum test with Wilcoxon Mann-Whitney U test were used to analysis statistically significant metabolites and genes between sample groups from the analysis of CRC tissues and normal colon tissue samples. Unpaired Student t-test was used to identify significant metabolites between MC38 cells lines with different treatments. All analysis were adjusted for multiple comparisons using Benjamini-Hochberg-based FDR correction. A two-sided adjusted p value ≤ 0.05 was considered statistically significant. Missing values of metabolite abundance were imputed using KNN (fold=5).

Initial sPLS-DA model was built with mixomics package (function “splsda”), the optimal number of components was selected by tuning, then the final sPLS-DA model was established. Finally, R^2^ and Q^2^ values (**Figure 2**) were acquired through a permutation test performed with 300 random permutations in a sPLS-DA model.

**Figure 2.**
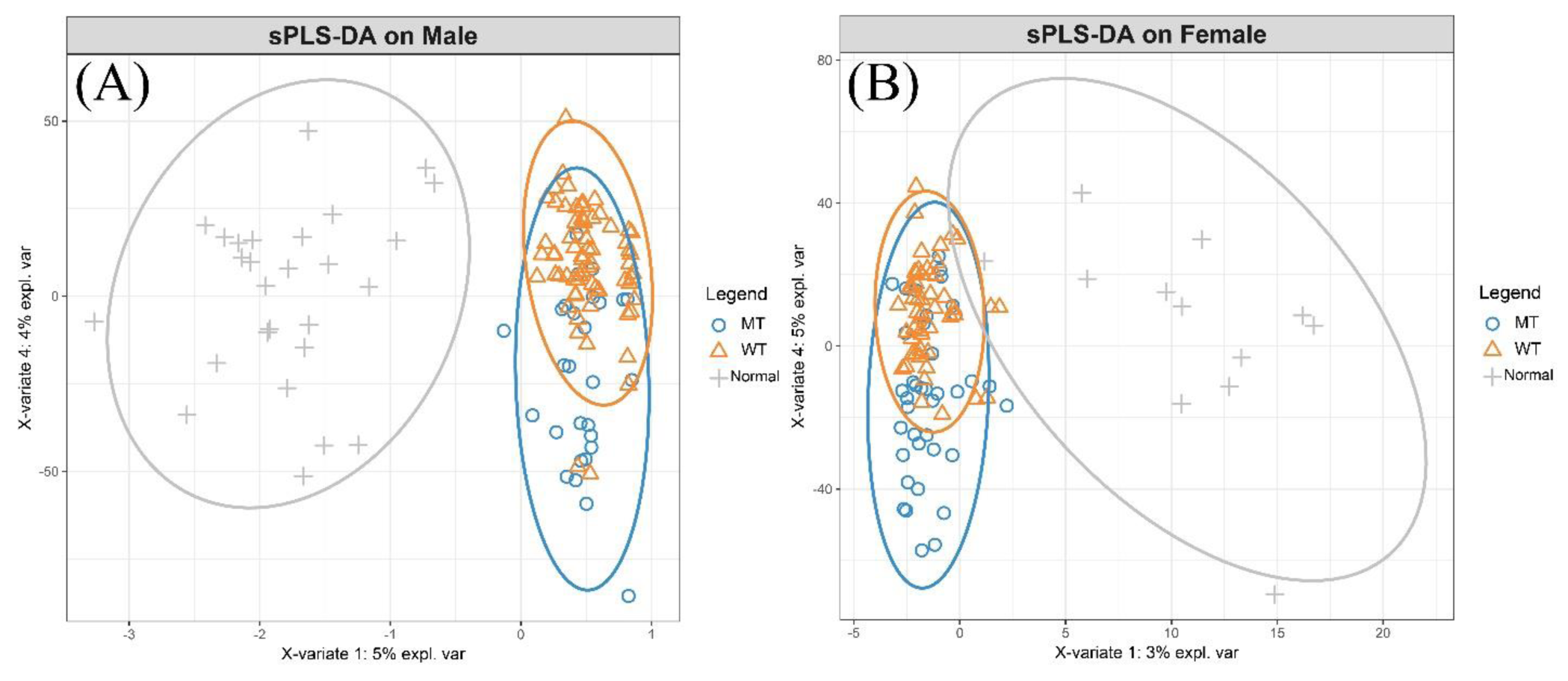
Metabolic differences in colon cancer revealed by untargeted metabolome profiling. Sparse partial least-squares discriminant analysis (sPLS-DA) scores plot assessing showing variance of metabolic features between normal colon tissues, KRAS mutant CRCs and KRAS wild-type CRCs. (A) Tumors and normal colon tissues from male patients: KRAS mutant CRCs (n=27), KRAS wild type CRCs (n=58), normal colon tissues (n=27); R2X (0.403), R2Y (0.98), Q2 (0.638), (B) Tumors and normal colon tissues from female patients: KRAS mutant CRCs (n=35), KRAS wild type CRCs (n=43), normal colon tissues (n=12); R2X (0.432), R2Y (0.987), Q2(0.753)

### Survival Analysis

Survival analyses on colon cancer patients (n = 397) from GSE39582 were carried out to investigate differences in outcomes between patients with KRAS mutant CRCs and patients with KRAS wild type CRCs by male or female sex. In addition to gene expression data, patient data including clinical parameters such as sex, age, survival status, and OS time were also downloaded. Patients with the colon cancer type colon adenocarcinoma (COAD) were selected. OS analyses were carried out only on patients with available survival and mRNA expression data. Two groups of samples: High GPX4 expression (with Z-Scores above upper quartile: 75%) and low GPX4 expression (with Z-Scores equal or below upper quartile: 25%) were categorized as previously described(9). OS analyses of FTH1, FTL and ACSL4 are the same as for GPX4. Kaplan-Meier survival analyses and Cox Proportional-Hazards models were conducted in R programming language using R packages “survminer” and “survival”. Log-rank test was utilized to analyze the statistical differences between different groups of colon cancer patients. Sex-specific differences were examined in multivariate Cox PH models by including an interaction between sex and ferroptosis-related genes. One prognostic outcome was considered: 5-year OS.

## Results

### Non-targeted metabolomics reveals differences in oxidative stress-related metabolites between tissues by KRAS status and sex

Non-targeted metabolomics analysis was previously performed on tumor tissues and normal colon tissues from CRC patients(6). We paired this metabolomics data with clinical data to classify and assess the metabolism of tumors by KRAS status. We performed sparse partial least-squares discriminant analysis (sPLS-DA), which revealed clear discrimination between the normal tissues and CRC tissues from both male and female patients, which was marginally better for males (based on visual inspection of sample clustering and the model statistics) (**Figures 2A-B**). Furthermore, there was additional discrimination between metabolomes from KRAS mutant and KRAS wild type tumors, suggesting different abundances of tumor metabolites exist based on KRAS status.

We further examined the metabolites that are driving these phenotypes. In our previous study, metabolite levels were compared between all normal colon and tumor tissues from male and females independently, revealing tumor-specific metabolites for both sexes(6). In this study, we compared these metabolites by sex and by KRAS status (mutant or wild type). Of note the metabolites we compared are represented in the following pathways: glycolysis, pentose phosphate pathway (PPP), amino acid metabolism, methionine cycle, TSP and glutathione (GSH) synthesis.

The initial analysis revealed differences in metabolites between the three tissue types (normal colon, KRAS wild type tumor, KRAS mutant tumors) (**Figures 2A-B**). S-adenosylmethionine (SAM), a common co-substrate involved in the transfer of methyl groups and metabolized within the TSP, was increased in both male and female tumors (both KRAS genotypes) compared to normal tissues. In males, SAM levels in KRAS mutant compared to wild type tumors trended towards being significantly higher (p=0.065), and the ratio of SAM/SAH was increased in males on comparison of KRAS mutant to normal tissues (**Figure 3A**). In females, the ratio of SAM/SAH was increased in both tumor types when comparing to normal tissues (**Figures 3B**). The levels of GSH were significantly upregulated in male patients with KRAS mutations compared to normal tissues, however the oxidized form of GSH, GSSG, was not altered between tissue types. We then examined the ratio of GSH/GSSG which was higher in tumors compared to normal tissues. This was different according to KRAS subtype and indicated that the conversion of GSH to GSSG is decreased in male tumors compared to normal tissues, and more so in KRAS mutant cases (**Table 1**, **Figure 3A**). In females, GSH was increased in both tumor types compared to normal tissues, but there were no differences in GSSG or the GSH/GSSG ratio between any of the tissue types (**Figures 3B**). These results suggests that the antioxidative capabilities of the cancer cells could differ between KRAS tumor types and by sex.

**Figure 3.**
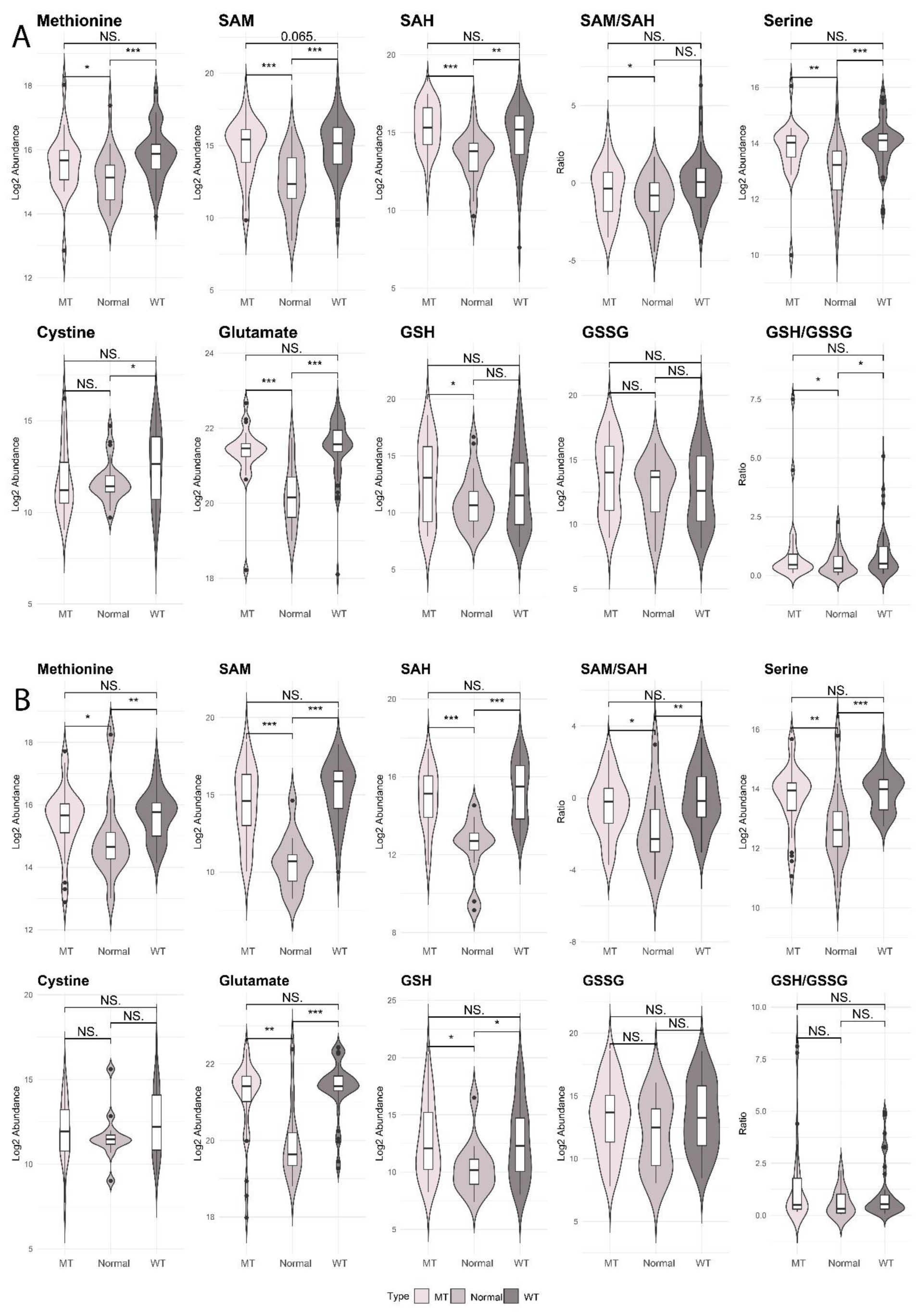
Critical metabolites required for methionine cycle, GSH biosynthesis and TSP in patients with CRC. A: Male: KRAS mutant CRCs (n=25), KRAS wild type CRCs (n=58), normal colon tissues (n=27); B: Female: KRAS mutant CRCS (n=35), KRAS wild type CRCs (n=43), normal colon tissues (n=12). Kruskal–Wallis rank sum test with Wilcoxon Mann-Whitney U test, p-values adjusted for false discovery rates (FDR) (Benjamini-Hochberg). *p < 0.05, **p < 0.01, ***p < 0.001, NS. = not significant. Abbreviations: GSH: reduced glutathione, GSSG: oxidized glutathione, SAM, S-Adenosylmethionine; SAH, S-Adenosylhomocysteine; TSP, transsulfuration pathway

### Semi-targeted analysis of metabolites related to ferroptotic pathways shows that KRAS mutant tumors exhibit decreased intratumoral ferroptosis in males only

As we found differences in oxidative stress-related pathways between tumor tissue KRAS subtypes and by sex, we performed a semi-targeted metabolomics analysis to examine additional metabolites nested in various pathways related to oxidative stress. To account for any metabolite degradation that may occur during longer term storage of tumor tissues that have already been extracted for metabolomics analysis, we re-extracted colon tissues and tumors from the same patient cohort and measured 29 metabolites in pathways related to the TSP, GSH metabolism, lipid peroxidation, mevalonate metabolism and inflammation. To confirm the identification and relative quantification of these metabolites, we compared commercially obtained metabolite standards to metabolites putatively identified in the metabolomics data (validated by measuring mass-to-charge ratio (*m/z*), retention time and MS/MS fragmentation patterns).

Comparing KRAS mutant to KRAS wild type tumors independent of sex, there are no significant metabolites altered in ferroptosis related pathways (**Supplementary Table 4**), however, when we split the group into female and male, there were two metabolites which were significantly decreased in abundance, and these differences were only seen in males: cystine, dihydrobiopterin (BH2) (**Table 1, Figure 4**). Additional metabolites in ferroptotic pathways were changed when comparing KRAS mutant or KRAS wild type tumors to normal colon tissues, in a sex-specific manner.

**Figure 4.**
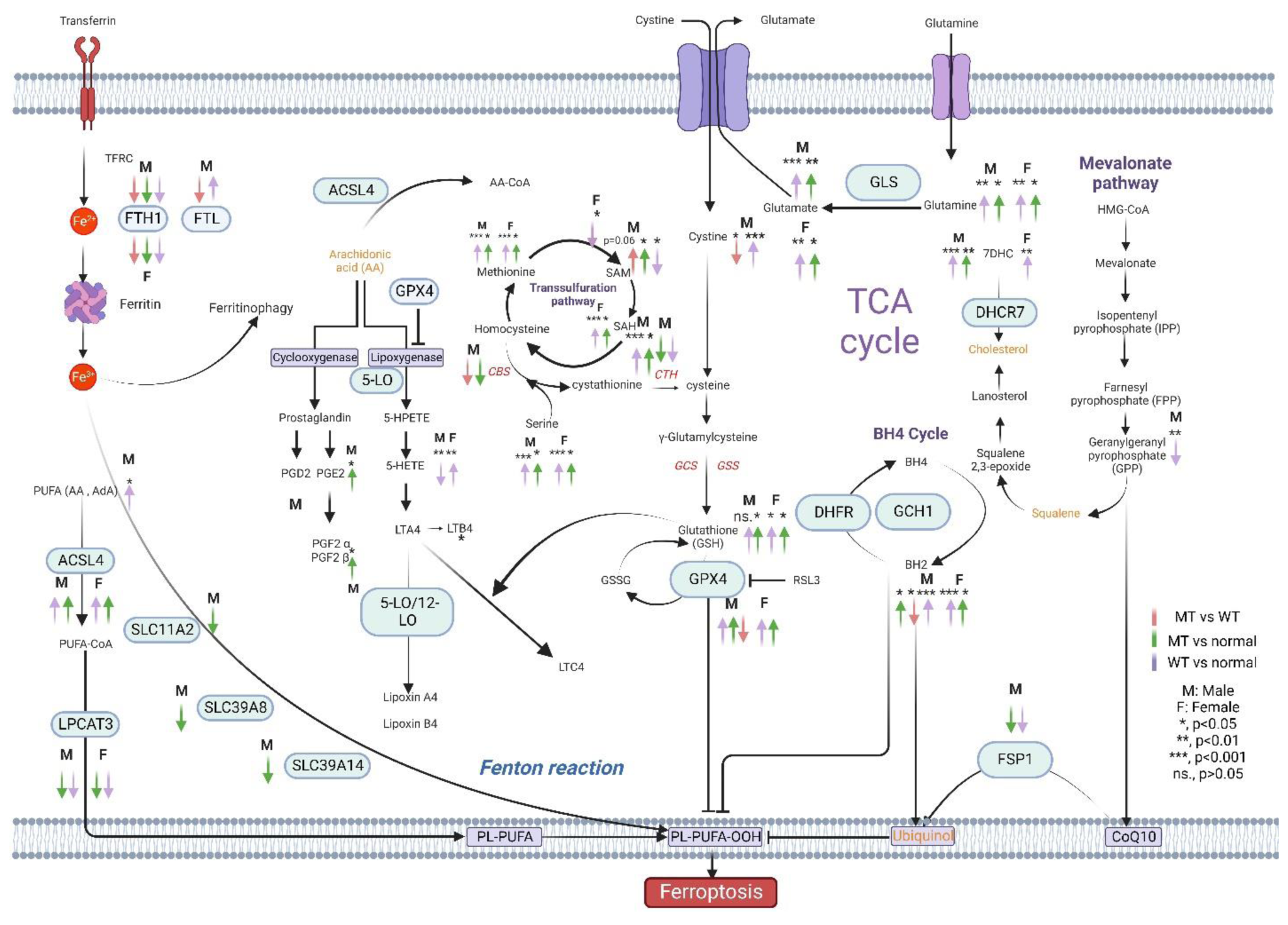
Scheme mapping changes to CRC-related metabolites within ferroptotic pathways. Arrows indicate directionality of change in metabolite abundance, color of arrow represents the tissue comparisons: red; KRAS Mutant CRCs compared to KRAS wild type CRCs, green; KRAS mutant CRCs compared to normal colon tissues, blue; KRAS wild type CRCs compared to normal colon tissues. Pathways shown describe the various mechanisms of ferroptosis: Cystine and cysteine availability, uptake through the xCT system or TSP pathway which is essential for GSH synthesis. The decreased accumulation of iron-dependent lipid peroxides by GPX4. The generation of substrates for lipid peroxidation (phospholipids with polyunsaturated acyl tails (PL-PUFAs) by enzymes such as ACSL4 and LPCATs (blue, bottom left) that activate and incorporate free PUFAs into phospholipids. Once PL-PUFAs are incorporated into membrane environments, iron-dependent enzymes and labile iron use molecular oxygen (O2) for peroxidation, generating PL-PUFA-OOH. There are three pathways for eliminating peroxidized PL-PUFAs (yellow, middle and bottom right): the GPX4-glutathione axis the FSP1-CoQ10 axis and the GCH1-BH4 axis. Iron-dependent enzymes including lipoxygenases and cytochrome P450 oxidoreductase (POR) also drive ferroptosis. Labile iron is imported through the transferrin receptor 1 (TfR1) and stored in ferritin. Ferritin can be degraded through an autophagy-like process known as ferritinophagy, which releases labile iron and facilitates the peroxidation reaction driving ferroptosis. **Abbreviations:** ACSL4, acyl coenzyme A synthetase long-chain family member 4; BH2, dihydrobiopterin; BH4, tetrahydrobiopterin; CoA, coenzyme A; Fe, iron; FSP1, ferroptosis suppressor protein 1; GCH1, GTP Cyclohydrolase 1; GCS, glutamylcysteine synthetase; GLS, glutaminase; GPX4, glutathione peroxidase 4; GSH: reduced glutathione, GSSG: oxidized glutathione, GSS, glutathione synthetase; HMG-CoA, 3-hydroxy-3-methylglutaryl CoA; LPCAT3, lysophosphatidylcholine acyltransferase 3; NRF2, nuclear factor E2-related factor 2; PE, piperazine erastin; PL, phospholipid; PUFA, polyunsaturated fatty acid; TCA, tricarboxylic acid; LPCAT, lysophosphatidylcholine acyltransferase; PUFA, polyunsaturated fatty acids, MUFA, Monounsaturated fatty acids; TFRC, Transferrin Receptor; PGD2, Prostaglandin D2; PGE2, Prostaglandin E2; PGF2α, Prostaglandin F2α; PGF2β, Prostaglandin FF2β; LTA4, Leukotriene A4; LTB4, Leukotriene B4; LTC4, Leukotriene C4; SAM, S-Adenosylmethionine; SAH, S-Adenosylhomocysteine; DHFR, Dihydrofolate reductase; DHCR7, 7-Dehydrocholesterol reductase; AA, Arachidonic acid; AdA: Adrenic acid

A closer investigation of cystine levels shows that in males, cystine was significantly increased in the KRAS wild type tumors compared to normal tissues and not different between KRAS mutant and normal tissues, suggesting differences in cystine between mutant and wild type may be driven by the increase in the KRAS wild type tumor levels. Cystine is used to generate GSH via cysteine and γ-glutamyl cysteine intermediates. GSH was increased only in KRAS mutant tumors compared to normal tissues. However, the GSH to GSSG ratio was significantly increased in both tumor types compared to normal tissues, suggesting that the conversion of GSH to GSSG by GPX4 was decreased in males. Given that GSH was only significantly upregulated in KRAS mutant tumors compared to normal tissues, this indicated that ferroptosis may be inhibited in male CRCs and more so in those with KRAS mutations. For female patients, the KRAS mutant and KRAS wild type tumors had similar changes in metabolites within this pathway compared to normal tumors. GSH were upregulated to a much higher level than for males, however neither GSSH nor the ratio of GSH to GSSG were altered.

Metabolites that contribute to the TSP have different trends in abundance between female and male CRCs based on KRAS status (**Table 1, Figure 4**). Methionine and SAH were upregulated in both KRAS wild type and KRAS mutant tumors compared to normal for both sexes. Whereas for SAM, this metabolite was upregulated in males, with KRAS mutant and downregulated in KRAS wild type compared to normal, however, SAM was only downregulated in KRAS wild type compared to normal in females. Further investigation of the ratio of SAM to SAH revealed a decreased ratio in KRAS mutant tumor compared to normal in both males and females, and a decreased ratio in KRAS wild type tumors compared to normal in females only. Other metabolites in this pathway were similarly changed in both sexes when comparing KRAS mutant and KRAS wild type tumors to normal tissues; serine which is a substrate for cystathionine production was increased in both KRAS types. Therefore, the metabolite levels in the TSP were changed in the same direction in all tumors compared to normal tissues for both males and females.

Examination of the tetrahydrobiopterin (BH4) cycle showed that in males, BH2 was decreased in KRAS mutant tumors compared to KRAS wild type tumors, this difference was not observed in females. However, the BH2 levels were upregulated in all tumors when compared to normal tissues for both sexes.

Ferroptosis is also associated with increased COX and LOX activity(10). These pathways are tightly connected to GPX4 and arachidonic acid (AA) metabolism, important molecules in ferroptosis(10) (**Table 1, Figure 4**). We examined AA levels; however, the levels of this metabolite were not different in abundance between tissue types in either sex. However, adrenic acid (AdA) another central intermediate in the LOX and COX pathway, was increased in KRAS wild type tumors compared to normal tissues for male patients only. Prostaglandins (PGs) PGE2 and PGF2β were also measured and significantly upregulated in KRAS mutant tumors compared to normal tissues in males. PGF2β, a downstream metabolite of PGE2, was also upregulated in KRAS mutant type tumors compared to normal in males. The PGs were not altered when comparing levels between tissues in females.

Studies have shown that increasing GPX4 activity can reduce pro-inflammatory lipid mediator production (i.e., 5-Hydroxyeicosatetraenoic acid (5-HETE)) driven by Nuclear factor kappa-light-chain-enhancer of activated B cells (NF-κB) pathway activation(11,12). In our data, we observed that 5-HETE was only decreased in males with KRAS wild type compared to normal tissues, conversely, in females, 5-HETE was increased in KRAS wild type compared to normal tissues. In the cholesterol synthesis pathway and mevalonate pathway, 7-dehydrocholesterol (7DHC) was increased in both KRAS mutant and KRAS wild type tumors in males, in females only dehydrocholesterol was increased in KRAS wild type tumors compared to normal tissues. Geranyl pyrophosphate (GPP) was only decreased in male KRAS wild type tumors compared to normal tissues (**Table 1, Figure 4**).

### Validation of metabolic phenotype for ferroptosis in KRAS mutant CRC cell lines

Given that our analysis of KRAS mutant tumors revealed potential ferroptosis suppression, we then sought to validate this metabolic phenotype in vitro, by activating ferroptosis in a CRC cell line. This was performed under the hypothesis that the metabolites altered with ferroptosis activation would be changed in the opposite direction or trend to those identified in the patient KRAS mutant tumors. We therefore treated KRAS mutant MC38 cells with RSL3, a ferroptosis inducer that inhibits GPX4, and performed semi-targeted metabolomics to identify ferroptosis metabolites on the harvested cell pellets. Changes to metabolite abundances involved in the ferroptosis pathway can be seen in **Supplementary Table 5** and **Supplementary Figure 2**. RSL3-treated cell lines had a trend of decreased GSH compared to MC38 cells treated with vehicle (i.e., controls), signifying alterations to GSH synthesis. GSSG which is a product of GSH metabolism by GPX4 was not altered, however the ratio of GSH/GSSG was decreased with RSL3 treatment. Similarly, in the TSP pathway, SAH was downregulated with RSL3 treatment and the ratio of SAM/SAH was upregulated. In addition, AA was upregulated. However, one metabolite in the COX pathway of AA metabolism was downregulated: PGE2. In addition to these pathways, metabolites in the BH4 cycle and mevalonate pathway were all downregulated in RSL3 treated cell lines: BH4, BH2, 7DHC and GPP. Thus, in our data set, it was evident that RSL3, a known GPX4 inhibitor and ferroptosis activator, downregulated GSH synthesis, the TSP, COX, and LOX pathways, the BH4 cycle and mevalonate pathway metabolism - all pathways that contribute to ferroptosis. The data from patients indicated that KRAS mutant tumors in male patients have decreased ferroptosis compared to KRAS wild type, a phenotype that was not observed in females. In males with KRAS mutant CRCs, many of the metabolites were dysregulated in an opposing direction compared to those altered in MC38 cells after RSL3 treatment, validating this finding.

### Validation of decreased ferroptosis in male patients with KRAS mutant CRC using gene expression data

To further validate our findings in CRC patients and KRAS mutant cell lines, we next examined differences in expression for genes that regulate ferroptosis pathways. We stratified by KRAS mutation status and patient sex using publicly available data from NCBIs GEO (GEO dataset GSE39582). **Tables 2** and **3** show the differential expression of ferroptosis-related genes between colon tumors by KRAS status and control colon tissues for males and females respectively.

**Table 2.** Differential expression of genes between tumor tissues by KRAS subtype and compared to normal tissues from male CRC patients using data from NCBI GEO: GSE39582. Wilcoxon Mann-Whitney U tests were performed, and p-values were adjusted for false discovery rates (Benjamini-Hochberg). The p-values and fold changes that are significantly altered are in bold. KRAS mutant tumors (n=82), KRAS wild type tumors (n=139), normal tissues (n=5)

| Male | adj.P value |  |  | Log2FC |  |  |
| --- | --- | --- | --- | --- | --- | --- |
|  | MT vs<br>WT | MT vs<br>normal | WT vs<br>normal | MT vs<br>WT | MT vs<br>normal | WT vs<br>normal |
| *Iron accumulation* |  |  |  |  |  |  |
| *NFE2L2* | 7.97E-01 | **4.02E-02** | **1.75E-03** | 0.05 | **-0.28** | **-0.35** |
| *FTH1* | **3.16E-03** | **3.38E-07** | **7.56E-07** | **-0.31** | **-1.83** | **-1.44** |
| *FTL* | **2.53E-02** | 4.82E-01 | **2.64E-03** | **-0.11** | 0.01 | **0.24** |
| *FECH* | 7.18E-01 | **1.25E-03** | **2.64E-03** | 0.05 | **-0.79** | **-0.73** |
| *ABCB6* | 8.48E-01 | **2.26E-03** | **2.31E-03** | 0.01 | **0.36** | **0.43** |
| *TFRC* | 8.45E-01 | 5.13E-01 | 4.43E-01 | 0.00 | 0.03 | -0.17 |
| *HMOX1* | 3.62E-01 | **8.95E-06** | **1.30E-03** | -0.17 | **-1.42** | **-1.02** |
| *NOX1* | 5.74E-01 | 4.16E-01 | **3.53E-02** | 0.26 | -0.50 | **-1.92** |
| *MT1G* | 7.53E-01 | **6.70E-06** | **1.14E-04** | -0.11 | **-2.20** | **-2.03** |
| *DPP4* | 3.95E-01 | **3.69E-02** | 1.92E-01 | 0.18 | **-0.82** | -0.38 |
| *NCOA4* | 3.62E-01 | **1.77E-02** | **2.43E-02** | -0.05 | **-0.58** | -0.53 |
| *Iron transport* |  |  |  |  |  |  |
| *SLC11A2* | 4.08E-01 | **3.73E-02** | 1.21E-01 | -0.13 | **-0.64** | -0.92 |
| *SLC39A14* | 6.01E-01 | **4.44E-02** | 6.06E-02 | -0.05 | **-0.61** | -1.17 |
| *SLC39A8* | 4.56E-01 | **3.91E-02** | 1.06E-01 | -0.11 | **-0.64** | -0.98 |
| *Lipid peroxidation* |  |  |  |  |  |  |
| *GPX4* | **7.96E-03** | **9.12E-04** | **5.92E-04** | **-0.23** | **0.41** | **0.76** |
| *AIFM2* | 7.13E-01 | **1.07E-03** | **5.04E-03** | -0.11 | **-0.86** | **-0.62** |
| *EGFR* | 5.52E-01 | 1.95E-01 | 3.14E-01 | -0.10 | -0.35 | -0.39 |
| *ACSL4* | 7.13E-01 | **8.20E-05** | **1.54E-02** | 0.09 | **1.14** | **1.04** |
| *LPCAT3* | 6.82E-01 | **1.23E-06** | **4.88E-05** | 0.05 | **-1.07** | **-0.86** |
| *ALOX12* | 8.03E-01 | **1.75E-02** | **5.04E-03** | 0.03 | **-0.23** | **-0.30** |
| *ALOX15* | 6.42E-01 | 9.08E-02 | 2.10E-01 | 0.01 | -0.22 | -0.22 |
| *GSH biosynthesis* |  |  |  |  |  |  |
| *SLC7A11* | 2.02E-01 | **4.14E-05** | **3.53E-02** | 0.29 | **1.94** | **0.74** |
| *SLC3A2* | 3.08E-01 | **2.74E-04** | **5.92E-04** | -0.14 | **0.56** | **0.68** |
| *GSS* | **1.40E-03** | 4.60E-01 | 3.46E-01 | **-0.27** | 0.02 | 0.13 |
| *GCLC* | 5.41E-01 | 4.04E-01 | 1.12E-01 | -0.09 | 0.06 | -0.27 |
| *GCLM* | 7.69E-01 | 2.56E-01 | 3.14E-01 | -0.08 | 0.23 | -0.22 |
| *NFS1* | **1.63E-02** | 9.19E-02 | **2.01E-03** | **-0.27** | 0.36 | **0.59** |
| *CDKN1A* | 5.83E-02 | **2.26E-03** | **1.13E-05** | 0.24 | **-1.00** | **-1.37** |
| *GSR* | 3.41E-01 | **4.04E-03** | **4.75E-03** | -0.10 | **-1.35** | **-1.26** |
| *Transsulfuration* |  |  |  |  |  |  |
| *pathway* |  |  |  |  |  |  |
| *CBS* | **2.18E-02** | **2.32E-02** | 2.42E-01 | **-0.28** | **-0.70** | -0.41 |
| *CTH* | 7.94E-01 | **5.15E-02** | **1.69E-04** | 0.03 | **-0.59** | **-1.42** |
| *Mevalonate pathway* |  |  |  |  |  |  |
| *HMGCR* | 7.20E-02 | 6.91E-01 | 4.13E-01 | 0.13 | -0.27 | -0.40 |
| *FDFT1* | 3.11E-01 | 2.05E-01 | 3.85E-01 | -0.05 | -0.32 | -0.27 |
| *Other* |  |  |  |  |  |  |
| *ATF4* | 8.02E-01 | **6.59E-05** | **4.66E-06** | 0.01 | **0.68** | **0.61** |

WT: KRAS wild type; MT: KRAS mutant; Adj.P value: p values adjusted for false discovery rates (FDR)

It can be seen from **Tables 2 and 3** that for both sexes *GPX4* was upregulated in both KRAS mutant and wild type tumors compared to normal colon tissues. In males only, *GPX4* is significantly decreased in KRAS mutant compared to KRAS wild type tumors. GPX4 oxidizes GSH to GSSG to prevent the accumulation of lipid peroxides and suppress ferroptosis. GSSG is then converted back to GSH via glutathione reductase (GR)-mediated reduction reaction, which consumes NADPH (**Figure 5)**. Since the expression or activity of GPX4 is controlled by GSH levels, the upregulation of *GPX4* in tumors may be a result of increased GSH synthesis and decreased inhibition of GPX4.

**Table 3.**
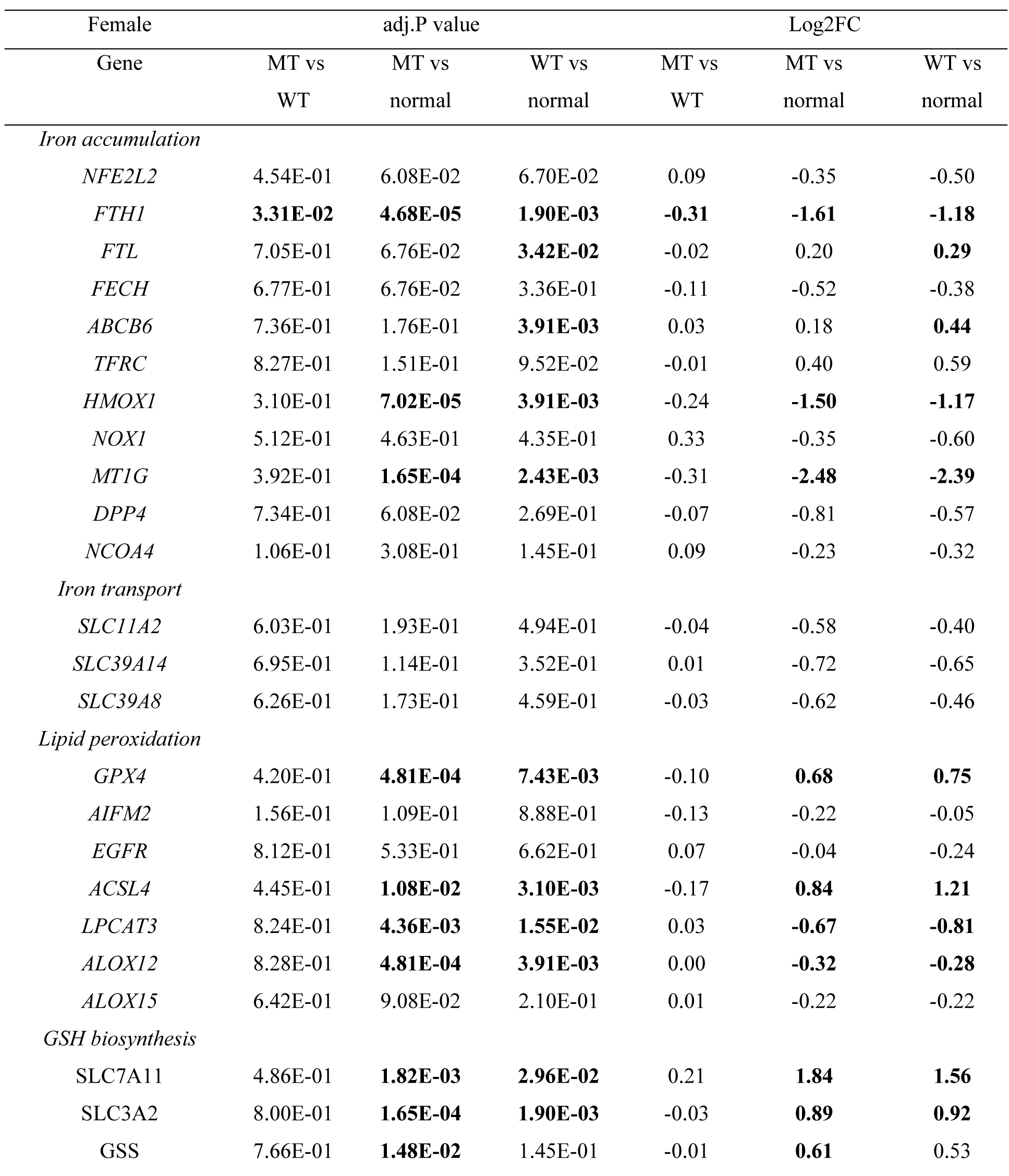
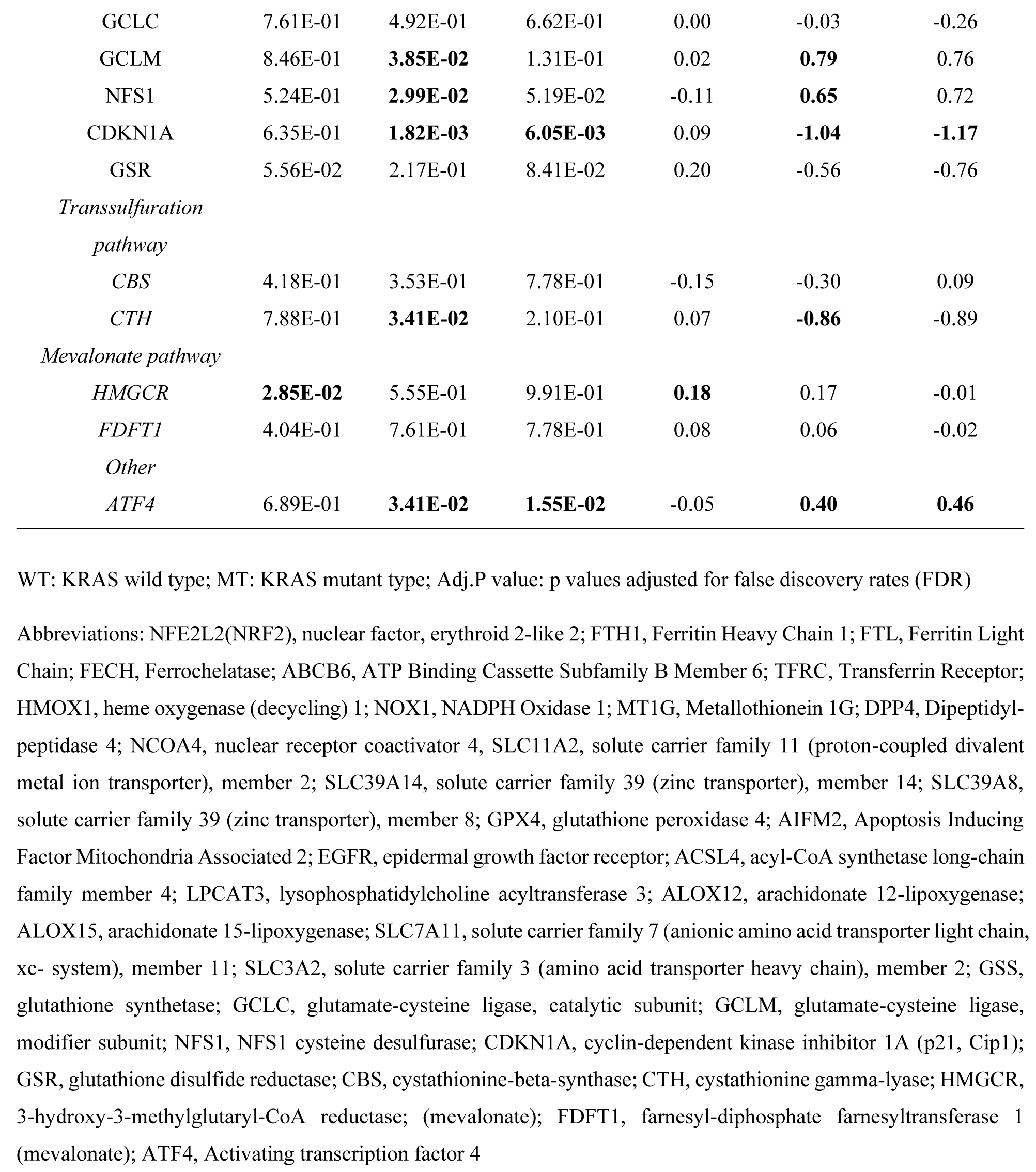
Differential expression of genes between tumor tissues by KRAS subtype and compared to normal tissues from female CRC patients using data from NCBI GEO: GSE39582. Wilcoxon Mann-Whitney U tests were performed, and p-values were adjusted for false discovery rates (Benjamini-Hochberg). The p-values and fold changes that are significantly altered are in bold. KRAS mutant tumors (n=66), KRAS wild type tumors (n=108), normal tissues (n=5)

| Female | adj.P value |  |  | Log2FC |  |  |
| --- | --- | --- | --- | --- | --- | --- |
| Gene | MT vs<br>WT | MT vs<br>normal | WT vs<br>normal | MT vs<br>WT | MT vs<br>normal | WT vs<br>normal |
| *Iron accumulation* |  |  |  |  |  |  |
| *NFE2L2* | 4.54E-01 | 6.08E-02 | 6.70E-02 | 0.09 | -0.35 | -0.50 |
| *FTH1* | **3.31E-02** | **4.68E-05** | **1.90E-03** | **-0.31** | **-1.61** | **-1.18** |
| *FTL* | 7.05E-01 | 6.76E-02 | **3.42E-02** | -0.02 | 0.20 | **0.29** |
| *FECH* | 6.77E-01 | 6.76E-02 | 3.36E-01 | -0.11 | -0.52 | -0.38 |
| *ABCB6* | 7.36E-01 | 1.76E-01 | **3.91E-03** | 0.03 | 0.18 | **0.44** |
| *TFRC* | 8.27E-01 | 1.51E-01 | 9.52E-02 | -0.01 | 0.40 | 0.59 |
| *HMOX1* | 3.10E-01 | **7.02E-05** | **3.91E-03** | -0.24 | **-1.50** | **-1.17** |
| *NOX1* | 5.12E-01 | 4.63E-01 | 4.35E-01 | 0.33 | -0.35 | -0.60 |
| *MT1G* | 3.92E-01 | **1.65E-04** | **2.43E-03** | -0.31 | **-2.48** | **-2.39** |
| *DPP4* | 7.34E-01 | 6.08E-02 | 2.69E-01 | -0.07 | -0.81 | -0.57 |
| *NCOA4* | 1.06E-01 | 3.08E-01 | 1.45E-01 | 0.09 | -0.23 | -0.32 |
| *Iron transport* |  |  |  |  |  |  |
| *SLC11A2* | 6.03E-01 | 1.93E-01 | 4.94E-01 | -0.04 | -0.58 | -0.40 |
| *SLC39A14* | 6.95E-01 | 1.14E-01 | 3.52E-01 | 0.01 | -0.72 | -0.65 |
| *SLC39A8* | 6.26E-01 | 1.73E-01 | 4.59E-01 | -0.03 | -0.62 | -0.46 |
| *Lipid peroxidation* |  |  |  |  |  |  |
| *GPX4* | 4.20E-01 | **4.81E-04** | **7.43E-03** | -0.10 | **0.68** | **0.75** |
| *AIFM2* | 1.56E-01 | 1.09E-01 | 8.88E-01 | -0.13 | -0.22 | -0.05 |
| *EGFR* | 8.12E-01 | 5.33E-01 | 6.62E-01 | 0.07 | -0.04 | -0.24 |
| *ACSL4* | 4.45E-01 | **1.08E-02** | **3.10E-03** | -0.17 | **0.84** | **1.21** |
| *LPCAT3* | 8.24E-01 | **4.36E-03** | **1.55E-02** | 0.03 | **-0.67** | **-0.81** |
| *ALOX12* | 8.28E-01 | **4.81E-04** | **3.91E-03** | 0.00 | **-0.32** | **-0.28** |
| *ALOX15* | 6.42E-01 | 9.08E-02 | 2.10E-01 | 0.01 | -0.22 | -0.22 |
| *GSH biosynthesis* |  |  |  |  |  |  |
| *SLC7A11* | 4.86E-01 | **1.82E-03** | **2.96E-02** | 0.21 | **1.84** | **1.56** |
| *SLC3A2* | 8.00E-01 | **1.65E-04** | **1.90E-03** | -0.03 | **0.89** | **0.92** |
| *GSS* | 7.66E-01 | **1.48E-02** | 1.45E-01 | -0.01 | **0.61** | 0.53 |
| GCLC | 7.61E-01 | 4.92E-01 | 6.62E-01 | 0.00 | -0.03 | -0.26 |
| GCLM | 8.46E-01 | **3.85E-02** | 1.31E-01 | 0.02 | **0.79** | 0.76 |
| NFS1 | 5.24E-01 | **2.99E-02** | 5.19E-02 | -0.11 | **0.65** | 0.72 |
| CDKN1A | 6.35E-01 | **1.82E-03** | **6.05E-03** | 0.09 | **-1.04** | **-1.17** |
| GSR | 5.56E-02 | 2.17E-01 | 8.41E-02 | 0.20 | -0.56 | -0.76 |
| *Transsulfuration* |  |  |  |  |  |  |
| *pathway* |  |  |  |  |  |  |
| CBS | 4.18E-01 | 3.53E-01 | 7.78E-01 | -0.15 | -0.30 | 0.09 |
| CTH | 7.88E-01 | **3.41E-02** | 2.10E-01 | 0.07 | **-0.86** | -0.89 |
| *Mevalonate pathway* |  |  |  |  |  |  |
| HMGCR | **2.85E-02** | 5.55E-01 | 9.91E-01 | **0.18** | 0.17 | -0.01 |
| FDFT1 | 4.04E-01 | 7.61E-01 | 7.78E-01 | 0.08 | 0.06 | -0.02 |
| *Other* |  |  |  |  |  |  |
| ATF4 | 6.89E-01 | **3.41E-02** | **1.55E-02** | -0.05 | **0.40** | **0.46** |

WT: KRAS wild type; MT: KRAS mutant type; Adj.P value: p values adjusted for false discovery rates (FDR)

**Figure 5.**
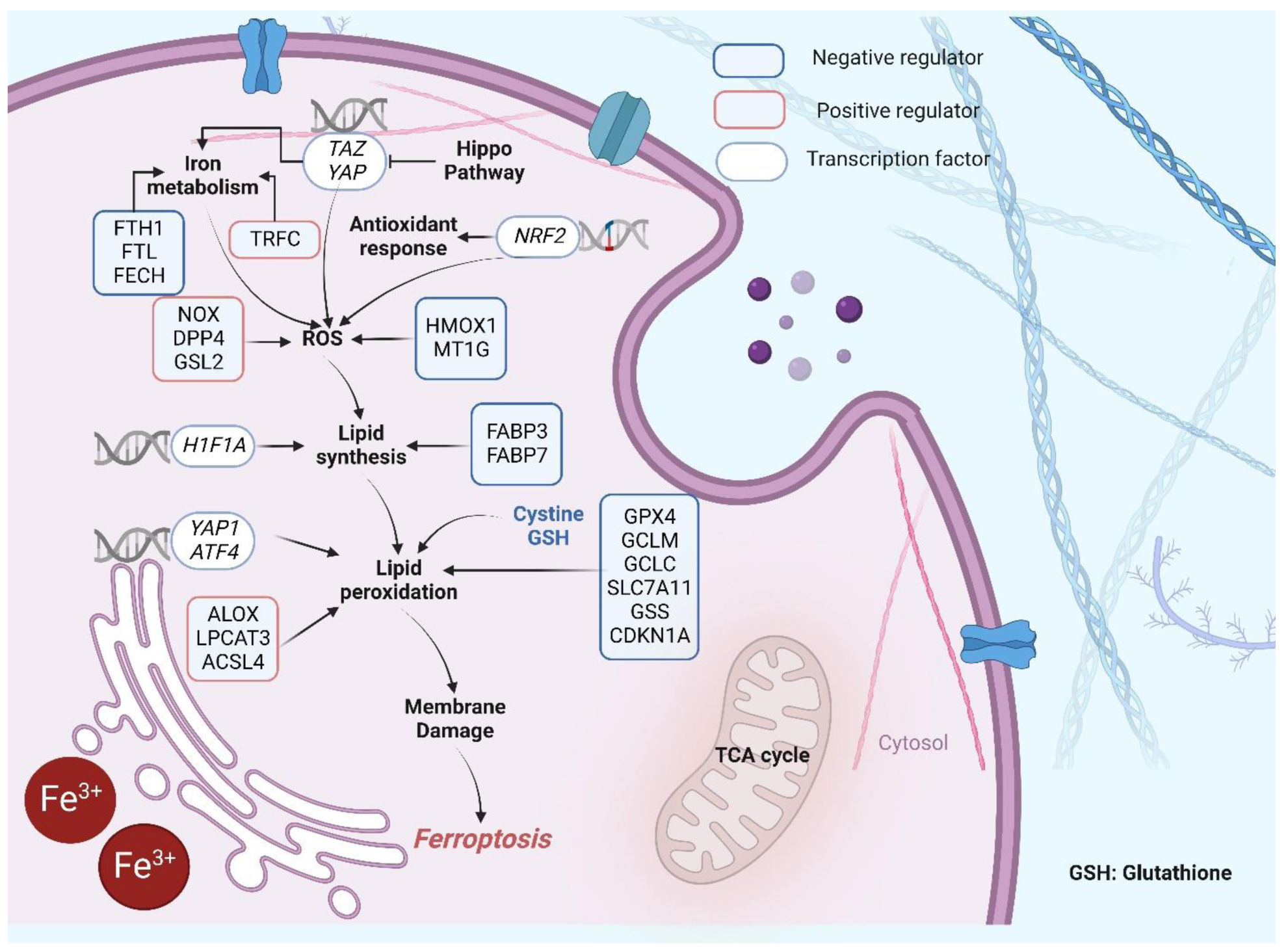
Key regulators of Ferroptosis. Ferroptosis is characterized by the production of lipid peroxidation. This process is positively regulated by iron-mediated reactive oxygen species production (NOX, DPP4, GSL2, TRFC), ACSL4-mediated lipid biosynthesis, and ALOX-mediated lipid peroxidation in cell membranes. On the contrary, the GSH pathway (regulated by GCLC, GPX4, GCLM, SLC7A11, GSS and CDKN1A), FABP3 and FABP7 (Upregulation of fatty-acid-binding protein 3 (FABP3) and fatty-acid-binding protein 7 (FABP7), which promotes fatty-acid uptake and lipid storage, driving the decline of ferroptosis (66)), play a parallel or complementary role in protection against membrane injury. YAP1 is transcriptional regulator in cell density mediated ferroptosis resistance, NFE2L2 maintains cell redox homeostasis by combining with an antioxidant response element in target genes. Abbreviations: TAZ, Transcriptional coactivator with PDZ-binding motif; YAP, Yes-associated protein 1; TRPC, Transferrin Receptor; FTH1, Ferritin Heavy Chain 1; FTL, Ferritin Light Chain; FECH, Ferrochelatase; NOX, NADPH Oxidase; DPP4, Dipeptidyl Peptidase 4; GSL2, Glutaminase 2; HIF1A, Hypoxia Inducible Factor 1 Subunit Alpha; YAP1, Yes1 Associated Transcriptional Regulator; ATF4, Activating Transcription Factor 4; ALOX, Arachidonate Lipoxygenase; LPCAT3, Lysophosphatidylcholine Acyltransferase 3; ACSL4, Acyl-CoA Synthetase Long Chain Family Member 4; HMOX1, Heme Oxygenase 1; MT1G, Metallothionein 1G; FABP3, Fatty Acid Binding Protein 3; FABP7, Fatty Acid Binding Protein 7; GPX4, Glutathione Peroxidase 4; GCLM, Glutamate-Cysteine Ligase Modifier Subunit; GCLC, Glutamate-Cysteine Ligase Catalytic Subunit; SLC7A11, Solute Carrier Family 7 Member 11; GSS, Glutathione Synthetase; CDKN1A, Cyclin Dependent Kinase Inhibitor 1A

GSH synthesis is mediated by both the TSP and xCT system. The TSP transfers sulfur from homocysteine to cysteine *via* cystathionine for biosynthesis of cysteine by cystathionine β-synthase (CBS) and cystathionine γ-lyase (CTH) (**Figure 4**)(13-15). CBS is allosterically regulated by SAM, which activates and increases the stability of CBS. Its activity increases flux through the TSP, which leads to GSH production. Comparison of gene expression in the TSP showed that in males, *CBS* expression was downregulated in KRAS mutant tumors compared to KRAS wild tumors, and in KRAS mutant compared to normal colon tissue groups. In addition, *CTH* was downregulated in both tumor subtypes compared to normal colon tissues, and the fold change decrease was greater in KRAS mutant tumors. This finding was not observed in female CRCs. On the contrary, *CBS* had no differential expression between tissue types, but *CTH* was downregulated in KRAS mutant tumors compared to normal tissues. This indicates decreased cysteine production in KRAS mutant tumors compared to normal tissues for both sexes, which was more pronounced in males when comparing to KRAS wild type tumors.

The xCT system functions as a cystine/glutamate transporter, that imports cystine in exchange for glutamate. This enables the generation of GSH via the conversion of cystine to cysteine. In both sexes, *SLC7A11*, a component of xCT system, was upregulated in KRAS mutant and KRAS wild type tumors compared to normal tissues. Notably, *SLC7A11* was upregulated to a greater extent in the KRAS mutant subgroup. *ATF4* is a positive regulator of *SLC7A11* under conditions of oxidative stress and cysteine deprivation(16). *ATF4* was also similarly upregulated. Of note, nuclear factor erythroid 2-related factor 2 (NFE2L2) is a master regulator of the antioxidant response and notably a regulator of *SLC7A11* via the antioxidant response element (ARE), present in the promoter region of the Xc-gene(17-19). In our analysis *NFE2L2* expression was downregulated in males only when comparing KRAS mutant and KRAS wild type tumors to normal colon tissues. Glutathione synthetase (GSS) and γ-glutamate-cysteine ligase (GCLC/GCLM) are also NFE2L2-target genes. GCLC and GCLM convert cysteine to γ-glutamyl-cysteine and their genes were not differentially expressed between tissue subgroups in males. *GSS*, which converts γ-glutamyl-cysteine to GSH, was downregulated in the male KRAS mutant group compared to KRAS wild type but was not different between tumor types and normal tissues. In females, *GSS* and *GCLM* were upregulated in KRAS mutant compared to normal tissues. Other genes involved in this pathway include cysteine desulfurase (*NFS1*) which was upregulated in KRAS wild type tumors compared in normal tissues in males and decreased in KRAS mutant tumors compared to wild type. In females, *NFS1* was increased KRAS mutant tumors compared to normal tissues only. Inhibition of *NFS1* has been reported to trigger ferroptosis. In addition, cyclin-dependent kinase inhibitor 1A (*CDKN1A*), which has been shown to inhibit ferroptosis(20,21), is decreased in both tumor subtypes compared to normal tissues in males, more so in KRAS mutant tumors, whereas in females is it decreased only in KRAS mutant compared to normal tissues. In conclusion, KRAS mutant tumors have decreased cysteine generation by the TSP and instead increase the import of cystine for generation of cysteine *via* the xCT, this was more prominent in males. In females, the xCT is activated in both KRAS mutant and wild type tumors, however cysteine catabolism to GSH is only increased in KRAS mutant tumors. Both KRAS mutant and wild type tumors have increased and similar *GPX4* activity compared to normal tissues which inhibits ferroptosis. In males, cysteine conversion to GSH does not appear to be active in either KRAS mutant or wild type tumors and is decreased more so in the KRAS mutant tumors compared to wild type. However, *GPX4* activity is increased in both tumor subtypes but is lower in the KRAS mutant composed to KRAS wild type tumors. GSH synthesis was increased in the male KRAS mutant subgroup through increased cystine transport and *GSS* downregulation, mediated to resist ferroptosis.

*ACSL4* esterifies CoA to free fatty acids and has a preference for long chain PUFAs such as AA and Ada(22). It is considered a specific biomarker and driver of ferroptosis, enhancing the PUFA content in phospholipids, which are susceptible to oxidation reactions leading to ferroptosis. In both sexes, *ACSL4* was upregulated in KRAS mutant and KRAS wild type tumors compared to normal tissues. However, lysophospholipid acyltransferase 5 *(LPCAT3)*, which is a positive regulator for lipid peroxide production and re-esterifies PUFA-CoAs into lysophospholipids, was downregulated in both KRAS mutant and wild type tumors compared to normal tissues in both sexes, showing resistance to ferroptosis. The arachidonate lipoxygenase *(ALOX)* genes, which play a dependent role in mediating PUFA peroxidation, were also examined. *ALOX12* was downregulated in KRAS mutant and wild type tumors compared to normal tissues in both sexes, however, *ALOX15* was not altered. This indicates that lipid peroxidation may be decreased in both tumor subtypes compared to normal tissues in both sexes. The *ALOX* family of genes may not be the only regulators of lipid peroxidation in ferroptosis, and it remains unknown if other lipoxygenases have a similar role in lipid peroxidation.

Additional pathways also show sex-differences. Ferroptosis Suppressor Protein 1 *(FSP1),* which prevents lipid oxidation and ferroptosis by reducing coenzyme Q10 (CoQ10) to ubiquinol-10, was downregulated in males when comparing both tumor subtypes to normal tissues but was not altered in females which is contradictory to our tumor findings.

Iron accumulation is another key hallmark of ferroptosis, and ferritin heavy chain 1 (*FTH1*) and ferritin light chain (*FTL*) are NFE2L2-target genes that mediate iron storage. *FTH1* was downregulated in KRAS mutant compared to KRAS wild type tumors, as well as in both tumor subtypes compared to normal tissues in males and females. In both sexes, *FTL* was increased in KRAS wild type tumors compared to normal but not changed in KRAS mutant tumors compared to normal. However, *FTL* was downregulated in the male KRAS mutant subgroup compared to wild type and was not altered when comparing these groups in females. We also examined the expression of the transferrin receptor (*TFRC*), which is regarded as a positive regulator for iron accumulation, however, *TFRC* expression was not significantly different between subgroups. In males, a number of additional genes related to iron accumulation were differentially expressed between tumors and normal tissues. Protoporphyrin/coproporphyrin ferrochelatase (*FECH)* was downregulated, and ATP-binding cassette, subfamily B (MDR/TAP), member 6 (*ABCB6)* was upregulated. In females, only *ABCB6* (**Figure 5**) was increased in KRAS wild type compared to normal tumors. The genes that link the mechanisms of iron accumulation to ROS generation were also altered in tumor subtypes. In males, negative regulators heme oxygenase 1 (*HMOX1*) (**Figure 5**) and metallothionein 1G (*MT1G*) (also NFE2L2 targets), were decreased in both subtypes of tumors compared to normal colons. Whereas positive regulators dipeptidyl-peptidase 4 (*DPP4*) and NADPH oxidase 1 (*NOX1*) were decreased in KRAS mutant tumors compared to normal tissues, and decreased in KRAS wild type tumors compared to normal tissues respectively. In females there were fewer genes altered, however, similarly to males *MT1G* and *HMOX1* were significantly decreased in tumors compared to normal tissues.

The uptake or transport of ferrous and ferric ions are also important for inducing ferroptosis. As illustrated in **Figure 4**, solute carrier family 39 (zinc transporter), member 8 (*SLC39A8), SLC39A14* and *SLC11A2* are transporters of ferrous ions. The expression of all three transporters was significantly decreased in KRAS mutant tumors compared to the normal colon tissues in males. These genes were not differentially expressed between any of the other subgroups or in females. This indicates that KRAS mutant CRCs from males have decreased ferrous ion transport, resulting in decreased lipid peroxidation. Overall, the results show that KRAS mutant tumors from male CRC patients have stronger capability for ferroptosis resistance.

### Sex-specific differences in the associations between ferroptosis-related genes and CRC prognosis according to KRAS status

Survival analysis was carried out to determine the association between sex, KRAS mutation and survival using gene expression data from the GSE39582 dataset. We show that male CRC patients with KRAS mutations have significantly poorer OS compared to those with KRAS wild type tumors (p=0.013) (**Figure 6A**). Strikingly, in female CRC patients there is no significant difference in OS between KRAS mutant and KRAS wild type tumors (p=0.34) (**Figure 6B**).

**Figure 6.**
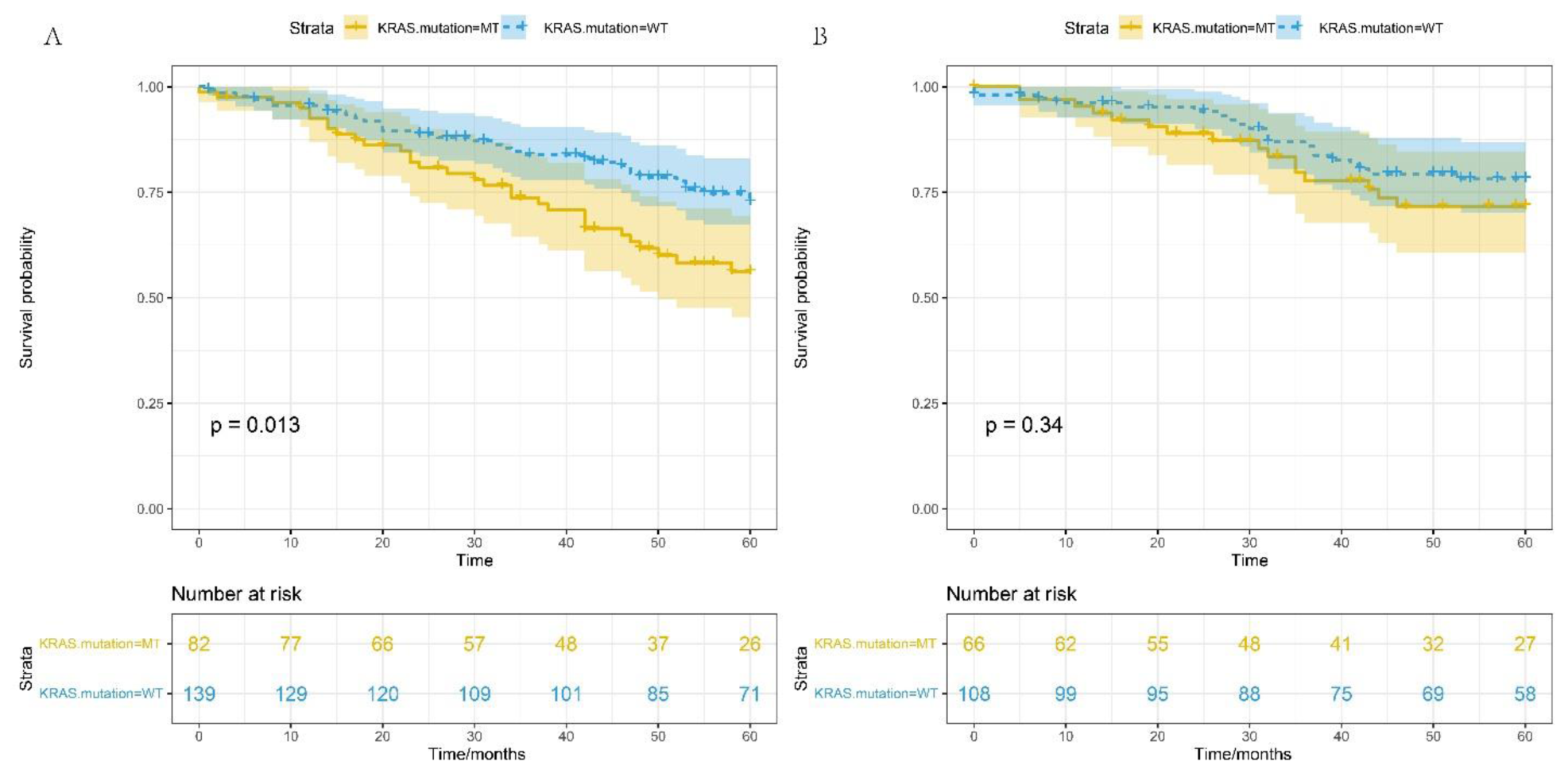
Cox proportional hazard survival curves for CRC patients with KRAS mutations and KRAS wild type tumors. A) OS for males, B), OS for females. Clinical data obtained from NCBIs GEO GSE39582 dataset, adjusted for anatomic location, chemotherapy history, clinical stage and age (continuous)

As this data indicates that male patients with KRAS mutations have poorer OS than those with KRAS wild type, we speculated that genes linked to the xCT-GSH-GPX4 axis and iron transport could also be associated with patient OS. According to our hypothesis, *GPX4* has higher expression in KRAS mutant type male CRCs compared to normal tissues which helps cancer cells grow by inhibiting ferroptosis. To determine if these patients would have poorer clinical outcomes we performed Kaplan-Meier survival analysis. **Figure 7** shows that high *GPX4* expression is associated with poorer OS in male CRCs with KRAS mutations (p = 0.011, log-rank test). However, *GPX4* is not associated with OS in males with KRAS wild type tumors. *GPX4* expression is not associated with OS in female CRC patients for those with KRAS mutant or KRAS wild type tumors.

**Figure 7.**
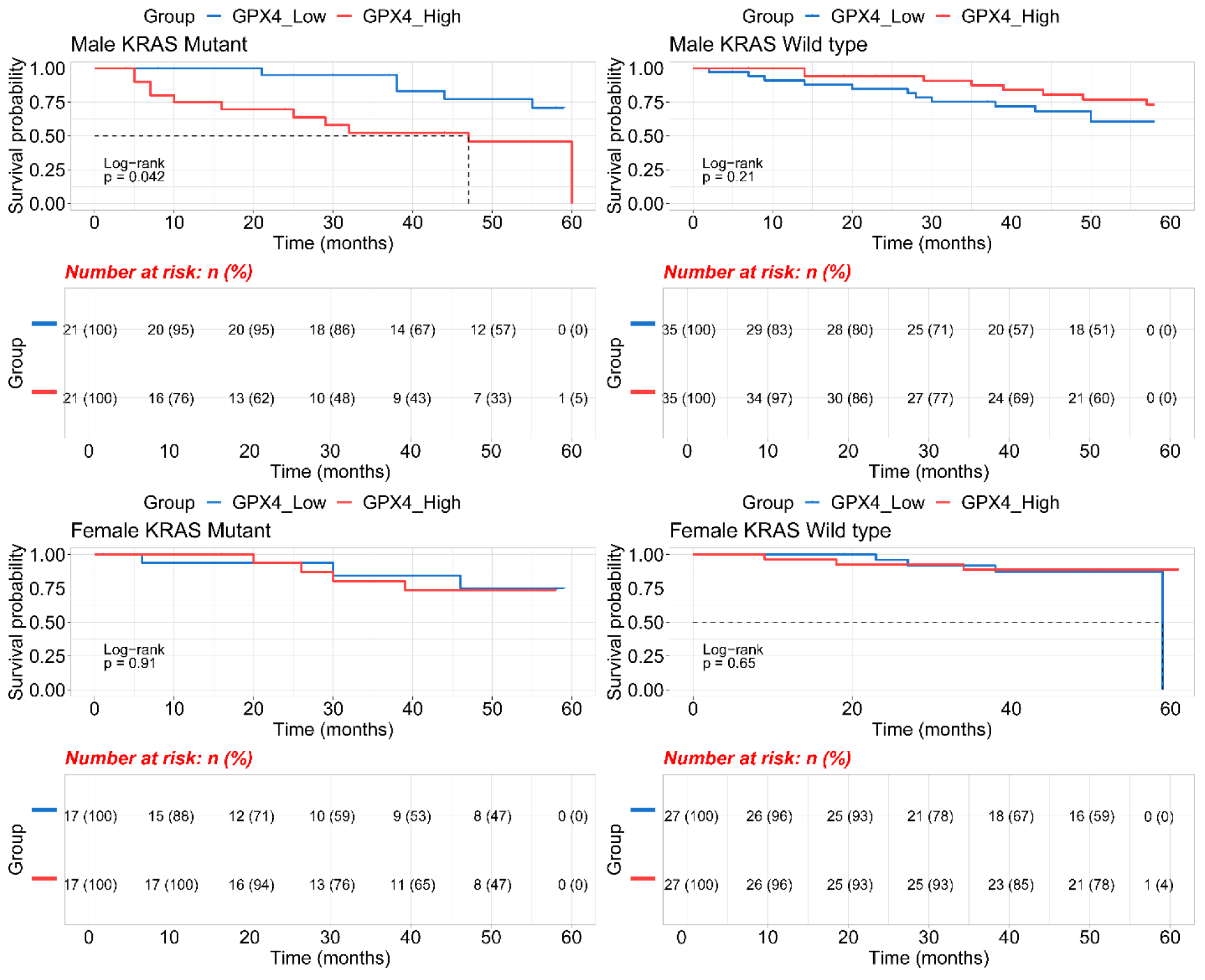
Kaplan–Meier survival curve and *GPX4* expression levels (low vs high) in CRC patients by sex and KRAS mutation type. Gene expression data obtained from NCBIs GEO GSE39582 dataset.

Further investigation of ferroptosis-related genes revealed significant associations between *FTH1*, *FTL* and *ACSL4* with OS for male and female CRCs stratified by KRAS status. The results showed a significant association between high *FTL* expression and poorer OS for male CRCs with KRAS mutant tumors (FTL: p = 0.013, **Supplementary Figure 3**). In addition, high *FTH1* expression trended towards an association with poorer OS for males with KRAS mutant CRC (p=0.056, **Supplementary Figure 4**), but there was no association between these genes and OS in male CRC patients with KRAS wild type tumors (FTL: p = 0.35, **Supplementary Figure 3**; FTH1: p=0.13, **Supplementary Figure 4**). For females, there was no association between these genes, and OS, similar to the findings observed for *GPX4*. In terms of *ACSL4*, a sex interaction based on KRAS status in the association between *ACSL4* and OS was observed. The results illustrated that low *ACSL4* expression trended towards poorer OS for male CRCs with KRAS mutations (p=0.056**, Supplementary Figure 5**), but no association in male CRCs with KRAS wild type and female CRCs (**Supplementary Figure 5**).

To examine whether these four genes were associated with OS after adjustment for anatomic location, clinical stage, and chemotherapy history, and age (continuous), we used cox proportional hazards multivariate models (**Supplementary Figure 6)**. Patients with KRAS wild type tumors had significantly better OS for males (HROS, 0.4; 95% CI, 0.23-0.69; p < 0.001) compared to males with KRAS mutant tumors. However, there was no evidence of an effect of KRAS status on female OS (HROS, 0.83; 95% CI, 0.40-1.7; p = 0.612). This data also shows that only high *FTH1* expression associates with worse OS for male CRC patients with KRAS mutations (HROS, 2.8; 95% CI, 1.21-6.63; p = 0.016) and was not associated with OS for females (HROS, 0.74; 95% CI, 0.28-1.9; p = 0.532).

These results validated our metabolomics data and shows a potential avenue for therapeutic intervention by regulating ferroptosis in male patients that have tumors with KRAS mutations and high *GPX4* expression.

## Discussion

In this study, we identified statistically significant sex-related differences in ferroptotic molecular pathways for CRC patients by KRAS subtype that associate with prognosis. KRAS mutant tumors from male CRC patients were altered in GSH biosynthesis, TSP activity and methionine metabolism. Additional validation from an independent dataset showed differential expression of genes involved in the regulation of these ferroptosis-related metabolic pathways. Examination of the relationship between these genes and survival by sex showed that only male patients with KRAS mutations have poorer 5-year OS compared to those with KRAS wild type tumors. Higher expression of *GPX4, FTH1, FTL* and lower *ACSL4* expression were also associated with poorer 5-year OS only in male patients with KRAS mutant tumors. The data also suggested that male KRAS mutant patients had increased iron dependence. Early onset colorectal cancer has a different biological process than older onset colorectal cancer, and thus we excluded this former subtype in our study.

It is known that cysteine is rate-limiting for the biosynthesis of GSH, and GSH levels directly correlate with cysteine availability. Therefore, a disruption of cysteine uptake, either genetically or chemically, efficiently depletes intracellular GSH levels(26). GSH, the most abundant reductant in mammalian cells enables the production of iron–sulfur clusters and is a substrate for multiple enzymes, including glutathione peroxidases (GPXs) and glutathione-S-transferases(27). In our study, GSH levels were upregulated in KRAS mutant tumors compared to normal tissues in males and females, however, they were only upregulated in KRAS wild type tumors compared to normal tissues in females. GSSG, the reduced form of GSH was not different in abundance between tissue types, but the ratio of GSH to GSSG was significantly higher in both KRAS mutant and KRAS wild type tumors compared to normal tissues for males only. This indicated that GPX4 could be inhibited more in tumors from males.

In addition, cystine, a precursor of cysteine imported by xCT was significantly decreased in KRAS mutant compared to KRAS wild type tumors from males, but this was not observed in females. We also examined the TSP in KRAS mutant CRCs compared to normal tissues. SAM was increased in males only, however SAH was increased in both sexes, further examination of the SAM/SAH showed a decreased ratio in KRAS mutant tumors from males and in both KRAS mutant and wild type tumors from females, enabling the provision of substrates for GSH in lieu of provision of cystine by xCT. Within the TSP, CBS enables the transfer of sulfur from homocysteine to cysteine via cystathionine, and recent evidence illustrated that the expression of *CBS* is inhibited by DNA methylation in tumors with KRAS mutations(31). The expression of *CBS* and cystathionase (*CSE*) are also dependent on the cancer subtype (i.e., KRAS mutant etc.). Given this finding we hypothesize that *CBS* is critical for TSP activity in KRAS mutant tumors from males. This hypothesis is validated by our findings from both the analysis of gene expression and metabolomics data for molecules within the TSP. When cysteine levels are low, the TSP cannot maintain GSH levels, therefore, the capacity for ROS scavenging will be decreased and ferroptosis triggered. Therefore, by combining the metabolomics and gene expression results, we observed that KRAS mutant tumors from males have suppressed ferroptosis through the system xCT-GSH-GXP4 pathway.

Iron dependence is another hallmark of ferroptosis. FTL and FTH1 store iron and can be degraded by lysosomes to increase free iron levels. Furthermore, the non-enzymatic, iron-dependent Fenton chain reaction is likely essential for ferroptosis: when GPX4 is inhibited, phospholipid hydroperoxides (PLOOHs) can persist longer, initiating the Fenton reaction to rapidly amplify PLOOHs. As the Fenton reaction is a significant driver process for PLOOH accumulation, ferrous and ferric ions are employed to generate the free radicals PLO• and PLOO• from PLOOH, respectively, driving the damaging peroxidation chain reaction. Our gene expression results suggest that free, redox-active iron levels were slightly increased in KRAS mutant tumors compared to KRAS wild type tumors in males only. In our results, the iron transporters *SLC11A2*, *SLC39A8*, and *SLC39A14* were downregulated in KRAS mutant tumors from males, therefore lipid peroxidation cannot occur via the Fenton reaction. Finally, survival analysis illustrates that higher expression of *GPX4*, *FTH1*, and *FTL* and lower expression of *ACSL4* were associated with poorer OS only in males with KRAS mutant CRC, which was consistent with our hypothesis that ferroptosis is decreased in these patient tumors. Our study suggests that these genes are likely more important drivers of KRAS mutant CRC in males compared to females that control CRC cell ferroptosis.

AA, an unsaturated fatty acid released from the cell membrane in response to various cytokines, peptides and growth factors(38), can be metabolized by COX, LOX, and cytochrome P450 (CYP450) monooxygenases to synthesize bioactive inflammatory mediators such as PGs, leukotrienes, epoxyeicosatrienoic acids and HETEs(39). In our data, we saw that PGs (PGE2 and PGF2β) were increased in KRAS mutant tumors from male patients only (compared to normal tissues), HETEs were not altered in KRAS mutant tissues from males or females, but they were decreased in KRAS wild type tumors compared to normal tissues in males and increased in females. These sex-difference show that inflammatory responses in male KRAS mutant group were decreased potentially due to decreased ferroptosis in these tumors.

As a potential antioxidant system, the BH4 cycle has also attracted attention. Although we only identified BH2 in our patient CRC datasets, we observed a significant difference in BH2 (oxidized metabolite from BH4) between mutant and wild type tumors driven by increased levels in the wild type KRAS tumors. In tumors from males with KRAS wild type, BH2 was increased compared to normal and KRAS mutant type. This indicated that tumors from males with KRAS wild type tumors might depend more on BH4 in the absence of GSH.

To determine whether the metabolic phenotype observed in the patient tumors is indicative of alterations to ferroptosis, we treated KRAS mutant MC38 CRC cell lines with RSL3, which induces ferroptosis. RSL3 directly inhibits GPX4, reducing antioxidant levels which leads to ferroptosis. In our dataset from CRC patients, GSH levels were upregulated in KRAS mutant tumors compared to normal tissues in males and females, however, it was only upregulated in KRAS wild type tumors compared to normal tissues in females. The ratio of GSH to GSSG was upregulated in both KRAS mutant and KRAS wild type tumors compared to normal tissues in males. On the contrary, this ratio was downregulated in MC38 cell lines. The ratio of SAM to SAH showed an opposite trend between KRAS mutant CRCs from males and MC38 cell lines. The ratio was upregulated in MC38 cell lines but downregulated in KRAS mutant type male CRCs. Furthermore, the AA downstream metabolite PGE2 had a reverse trend between patient tumors from males with KRAS mutations and MC-38 cells treated with RSL3, indicating that these tumors have decreased ferroptosis. This was not observed in females. As RSL3 was used to induce ferroptosis and a reverse trend of metabolite abundances in pathways linked to ferroptosis (TSP, GSH synthesis, COX and LOX) were observed in KRAS mutant tumors from males, it is evident that ferroptosis is decreased in these tumors.

KRAS mutations, which frequently occur in CRC, are the most difficult to target. There is increasing evidence that KRAS also mediates ferroptosis, inflammation and crosstalk with tumor immunity(43). KRAS may sustain high iron levels for CRC growth by regulation of HIF2α and JAK-STAT signaling, and HIF2α-mediated iron import through the divalent metal transporter 1 (DMT1)(44,45). Analysis of TCGA data has previously revealed that KRAS mutations potentiate the expression of iron importers in CRC(46). In KRAS-mutant pancreatic ductal adenocarcinoma (PDAC), autophagy-dependent ferroptosis was shown to promote the release of oncogenic KRAS to polarize macrophages toward an immunosuppressive phenotype(47). Also in PDAC, KRAS mutations were associated with increased redox capability via the conversion of glutamine to oxaloacetate, malate, and pyruvate(48). Furthermore, KRAS mutant cancer cell lines were found to be sensitive to ferroptosis inducers. Yet, it is still unclear what mechanism is used in mutant KRAS CRCs to regulates the levels of ferroptosis metabolites and genes to resist this type of cell death.

The competitive strengths of this study include in-depth analysis of the sex-differences underlying the metabolic profiles of CRC tumors based on KRAS status through untargeted and targeted metabolomics. Additionally, treatment of KRAS mutant cells with a ferroptosis inducer confirms that the metabolic phenotype observed in male patients. Gene expression and survival analysis from two additional data resources of CRC patients validates our hypothesis and indicates that decreased ferroptosis in male patients with mutant KRAS associates with poorer outcomes. Undeniably, more work is warranted to understand how KRAS alters ferroptosis metabolism in a sex-specific manner. These mechanisms will be important to understand to develop new therapies to target KRAS in a sex-specific manner.

## Additional Information

**Financial Support:** This research was funded by NIH 1R21CA223686-01 (C.H.J., S.A.K.) and American Cancer Society research scholar grant 134273-RSG-20-065-01-TBE (C.H.J.). C.H.J would also like to acknowledge support from the National Cancer Institute of the NIH under Award Number K12CA215110, the NIH/NCI Yale SPORE in Skin Cancer under award number 5P50CA121974. RT is supported by NIH/NCI F30CA254246. This publication was also made possible by CTSA grant number UL1 TR001863 from the National Center for Advancing Translational Science (NCATS), components of the National Institutes of Health (NIH), NIH roadmap for Medical Research, and Women’s Health Research at Yale. Its contents are solely the responsibility of the authors and do not necessarily represent the official view of NIH. This work was also supported by the Lampman Research Fund in Yale Surgical Oncology. R.T. is supported by NIH F30CA254246.

## Translational Relevance Statement

CRC represents a major cause of morbidity and mortality in both male and female patients. Approximately 40% of patients with CRC carry a KRAS mutation. Additionally, KRAS mutations in CRC are associated with tumor aggressiveness and are difficult to treat. In this study, we examined factors that affect survival in an additional cohort of patients with stage I-III CRC. We show that patients with KRAS mutant tumors have a poorer 5-year overall survival compared to those with KRAS wild type tumors, and this association only exists in male CRC patients. In addition, high expression of *GPX4, FTH1,* and *FTL* and low *ACSL4* expression were associated with poorer 5-year overall survival only in KRAS mutant tumors from male patients, and sex-specific metabolic signatures indicated a role of decreased ferroptosis in male CRC patients by KRAS status, revealing a potential avenue for therapeutic approaches.

## Notes

### Competing Interest Statement

The authors have declared no competing interest.

