## Supplemental tables and figures for "Discovery of decreased ferroptosis in male colorectal cancer patients with KRAS mutations"

***Abbreviations***

KRAS, Kirsten rat sarcoma viral oncogene homolog

RCC, right-sided colorectal cancer;

LCC, left-sided colorectal cancer;

MT, mutant type;

WT, wild type;

MPST, 3-mercaptopyruvate sulfurtransferase;

ATF4, activating transcription factor 4; CDO, cysteine dioxygenase; CSE,

SAM, S-adenosyl-methionine;

SAH, S-adenosyl-homocysteine;

r-Glu-Cys, γ-L-Glutamyl-L-cysteine

5-HETE, 5-Hydroxyeicosatetraenoic acid

GPP, Geranyl pyrophosphate

GSSG, Glutathione disulfide

4-HpNE, 4-hydroperoxy 2-Nonenal

4-HNE, 4-Hydroxynonenal

15S-HETE, 15-Hydroxyeicosatetraenoic acid

PGPC, 1-palmitoyl-2-glutaryl phosphatidylcholine

15(S)-HpETE-SAPC, 1-Stearoyl-2-15(S)-HpETE-sn-glycero-3-Phosphatidylcholine

TSP, transsulfuration pathway;

MPST, 3-mercapto-pyruvate sulfurtransferase;

**Sample collection**

Colon tumor and normal colon tissue was acquired from surgery and prospectively collected on 736 stage I-IV CRC patients in the period 1991–2001 at Memorial Sloan-Kettering Cancer Center (MSKCC, New York, NY, United States). Clinical data was recorded and updated retrospectively. Tumor tissue and normal colon tissue (away from the tumor at the resection margin) was acquired from surgical colectomy specimens. Each sample was snap frozen in liquid nitrogen and immediately stored in a −80 °C freezer. Pre-operative intravenous antibiotics (cefazolin/metronidazole, clindamycin/gentamicin or ciprofloxacin/metronidazole) were administered within 60 min prior to resection. All patients received a standard mechanical bowel preparation (polyethylene glycol (PEG) solution) 24 h before scheduled surgery. For this study, samples were selected from patients that were $\geqslant$55 years old to reduce the confounding effects of estrogen signaling on metabolism before menopause. All normal colon tissues were selected from stage I-IV CRC patients (n = 39), and tumor tissue samples were selected from RCCs and LCCs stage I-III (n = 197). Stage IV tumor samples were not included as their metabolism may be affected by the presence of metastases in the liver or other site, therefore we cannot rule out this confounder. Normal samples were taken from stage IV patients, as their metabolic profiles were not significantly different from the normal colon tissues taken from stage I-III patients (Supplementary Fig. 6), and were markedly different from the tumor sample metabolomic profiles. The Yale University IRB determined that the study conducted in this publication was not considered Human Subjects Research and did not require IRB review (IRB/HSC# 1612018746). The study does not obtain data through intervention or interaction with the individual, or does not use or obtain identifiable private information. Informed consent was waived as part of the study exemption

**UPLC-MS analysis**

Both HILIC-MS and RPLC-MS approaches were used for comprehensive analysis of the tissue metabolome. A UPLC system (H-Class ACQUITY, Waters Corporation, MA, United States) coupled to a quadrupole time-of flight (QTOF) mass spectrometer (Xevo G2-XS QTOF, Waters Corporation, MA, United States) was used for MS data acquisition. A Waters ACQUITY UPLC BEH Amide column (particle size, 1.7 μm; 100 mm (length) × 2.1 mm (i.d.)) and Waters ACQUITY UPLC BEH C18 column (particle size, 1.7 μm; 50 mm (length) × 2.1 mm (i.d.)) were used for the UPLC-based separation of metabolites. The column temperature was kept at 25 °C for HILIC-MS analysis and 30 °C for RPLC-MS analysis. The solvent flow rate was 0.5 mL/min, and the sample injection volume was 1 μL. For HILIC-MS analysis, mobile phase A was 25 mM NH4OH and 25 mM NH4OAc in water, while the mobile phase B was ACN for both electrospray ionization (ESI) positive and negative mode, respectively. The linear gradient was set as follows: 0~0.5 min: 95% B; 0.5~7 min: 95% B to 65% B; 7~8 min: 65% B to 40% B; 8~9 min: 40% B; 9~9.1 min: 40% B to 95% B; 9.1~12 min: 95% B. For RPLC-MS analysis, the mobile phases A was 0.1% formic acid in H2O, while the mobile phases B was 0.1% formic acid in ACN, respectively for both ESI+ and ESI−. The linear gradient was set as follows: 0~1 min: 1% B; 1~8 min: 1% B to 100% B; 8~10 min: 100% B; 10~10.1 min: 100% B to 1% B; 10.1~12 min: 1% B. Pooled samples were analyzed every eight injections during the UPLC-MS analysis to monitor the stability of the data acquisition and used for subsequent data normalization.

QTOF-MS scan data (300 ms/scan; mass scan range 50–1000 Da) was initially acquired for each biological sample for metabolite quantification. Then, both DDA (data-dependent acquisition) data (QTOF MS scan time: 50 ms/scan, MSMS scan time 50 ms/scan, collision energy 20 eV, top 5 most intense ions were selected for fragmentation, exclude former target ions (4 s after 2 occurrences)) and MSe data (low energy scan: 200 ms/scan, collision energy 6 eV; high energy scan: 100 ms/scan, collision energy 20 eV, mass scan range 25–1000 Da) were acquired for QC samples to enable metabolite identification. ESI source parameters on the Xevo GS-XS QTOF were set as following: capillary voltage 1.8 kV, sampling cone 40 V, source temperature 50 °C, desolvation temperature 550 °C, cone gas flow 40 L/Hr, desolvation gas flow 900 L/Hr.

MS/MS method: Collision energy ramp: 10~40V, all the metabolites were match with standards, including m/z, retention time and MS/MS.

**UPLC-MS data processing**

The raw MS data (.raw) were converted to mzML files using ProteoWizard MSConvert43 (http://proteowizard.sourceforge.net/). The files were then processed in R (version 4.0.3) using the XCMS package for feature detection, retention time correction and alignment. The XCMS processing parameters were optimized and set as follows: mass accuracy for peak detection = 20 ppm; peak width c = (5, 30); snthresh = 6; bw = 10; mzwid = 0.015; minfrac = 0.5. The CAMERA package was used for subsequent peak annotation. The resulting data were normalized using the support vector regression algorithm (MetCleaning) in R to remove unwanted system error that occurred among intra- and inter- batches.

### *Supplementary Tables*

**Supplementary Table 1.** Demographics of colorectal cancer patients from samples used for non-targeted metabolomics analysis.

| | **Normal (n=39)** | **KRAS mutant (n=60)** | **KRAS wild type (n=101)** |
|---|---|---|---|
| **Sex, n** | | | |
| Male | 27 | 25 | 58 |
| Female | 12 | 35 | 43 |
| **Age, mean (SD)** | | | |
| Male | 69.3 (9.6) | 71.2(7.3) | 70.1(8.6) |
| Female | 63.3 (16.1) | 70.3(7.5) | 73.5 (9.8) |
| **Race/Ethnicity, n** | | | |
| NHWs | 35 | 54 | 86 |
| Hispanic | 2 | 4 | 8 |
| AA | 1 | 1 | 4 |
| API | 1 | 1 | 3 |

NHWs: non-Hispanic White, AA: African-American, API: Asian-Pacific Islander, SD: Standard Deviation

**Supplementary Table *2*.** Summary of KRAS mutation type for colorectal cancer patient tumors analyzed by non-targeted metabolomics analysis.

| KRAS mutation | G12D | G12V | G12S | G12A | G12C | G13D |
|---|---|---|---|---|---|---|
| Male (n) | 8 | 9 | 1 | 3 | 1 | 3 |
| Female (n) | 17 | 10 | 1 | 0 | 1 | 3 |

**Supplementary Table 3.** Sex and tumor KRAS mutation status of patients from the NCBI GSE39582 dataset

| Patient Sex | KRAS mutant (n=148) | KRAS wild type (n=247) | non-tumor (n=10) |
|---|---|---|---|
| Female | 66 | 108 | 5 |
| Male | 82 | 139 | 5 |

**Supplementary Table 4.** Differences in metabolite levels between KRAS mutant and KRAS wild type tumors from CRC patients independent of sex. Wilcoxon Mann-Whitney U tests were performed, and p-values were adjusted for false discovery rate (Benjamini-Hochberg). KRAS mutant (n=70), KRAS wild type (n=101)

| | adj.P value | Log2 fold change |
|---|---|---|
| Metabolites | KRAS mutant VS KRAS wild type | KRAS mutant VS KRAS wild type |
| | ***xCT system*** | |
| Cystine | 3.34E-01 | 0.71 |
| Glutamine | 4.50E-01 | -0.10 |
| Glutamate | 3.17E-01 | 0.18 |
| GSH | 5.15E-01 | -0.24 |
| GSSG | 5.68E-01 | 0.00 |
| GSH/GSSG ratio | 7.60E-01 | 0.06 |
| | ***TSP and BH4 metabolism*** | |
| SAH | 9.93E-01 | 1.08 |
| SAM | 4.40E-01 | 0.26 |
| SAM/SAH ratio | 8.20E-01 | 0.12 |
| Methionine | 4.27E-01 | 0.14 |
| Serine | 4.40E-01 | 0.12 |
| Dihydrobiopterin | 6.40E-01 | -0.04 |
| | ***LOX and COX pathways*** | |
| Adrenic acid | 9.90E-01 | 0.12 |
| 5-HETE | 0.440430321 | 0.15 |
| Prostaglandin E2 | 7.92E-01 | 0.01 |
| Prostaglandin F2 β | 5.68E-01 | 0.15 |
| Arachidonic acid | 8.96E-01 | -0.04 |
| | ***Mevalonate and Cholesterol metabolism*** | |
| GPP | 6.82E-01 | 0.06 |
| 7-Dehydrocholesterol | 7.92E-01 | -0.07 |

Abbreviations: GSH: Glutathione; GSSG, Glutathione disulfide; SAM, S-adenosyl-methionine; SAH, S-adenosyl-homocysteine; 5-HETE, 5-Hydroxyeicosatetraenoic acid; GPP, Geranyl pyrophosphate; TSP, Transsulfuration pathway; BH4, Tetrahydrobiopterin

**Supplementary Table 5.** Changes to metabolites in MC-38 cells after treatment with ferroptosis inducer RSL3. Metabolites in bold are significantly altered after RSL3 treatment compared to control (vehicle). Paired student's t test was applied. p values adjusted for false discovery rates (FDR) (Benjamini-Hochberg). RSL3 treated (n=5), control (n=5)

| Metabolites | adj.P value | Log2 fold change |
|---|---|---|
| Comparison | RSL3 vs. control | RSL3 vs. control |
| ***xCT system*** | | |
| Cystine | 4.87E-01 | -1.51 |
| Glutamine | **1.15E-05** | **-1.57** |
| Glutamate | **3.99E-08** | **-4.29** |
| GSH | 8.52E-02 | -1.12 |
| GSSG | 1.72E-01 | 0.59 |
| GSH/GSSG | **5.91E-03** | **-1.85** |
| ***TSP and BH4 metabolism*** | | |
| SAH | **3.89E-05** | **-4.3** |
| SAM | 4.43E-01 | 0.4 |
| SAM/SAH | **2.71E-02** | **1.09** |
| Methionine | 5.43E-01 | -0.2 |
| Serine | **2.98E-05** | **-2.17** |
| BH4 | **1.41E-03** | **-1.27** |
| Dihydrobiopterin | **1.49E-03** | **-1.03** |
| ***LOX and COX pathways*** | | |
| Arachidonic Acid | **8.37E-03** | **2.28** |
| Adrenic acid | **4.36E-04** | **3.84** |
| 5-HETE | **6.87E-04** | **1.36** |
| Prostaglandin E2 | **4.65E-05** | **-1.99** |
| ***Mevalonate and Cholesterol metabolism*** | | |
| GPP | **4.17E-06** | **-3.03** |
| 7-dehydrocholesterol | **9.27E-03** | **-5** |

Abbreviations: GSH: Glutathione; GSSG, Glutathione disulfide; SAM, S-adenosyl-methionine; SAH, S-adenosyl-homocysteine; 5-HETE, 5-Hydroxyeicosatetraenoic acid; GPP, Geranyl pyrophosphate; TSP, Transsulfuration pathway; BH4, Tetrahydrobiopterin

**Supplementary Figure 1.** A) RSL3 dose titration, B) 10 μM RSL3 rescue with 100 μM ferrostatin-1, C) cellular lipid peroxide estimation with 10 μM RSL3 dose**.** ** $p < 0.01$, **** $p < 0.0001$ as determined by two-tailed t-test.

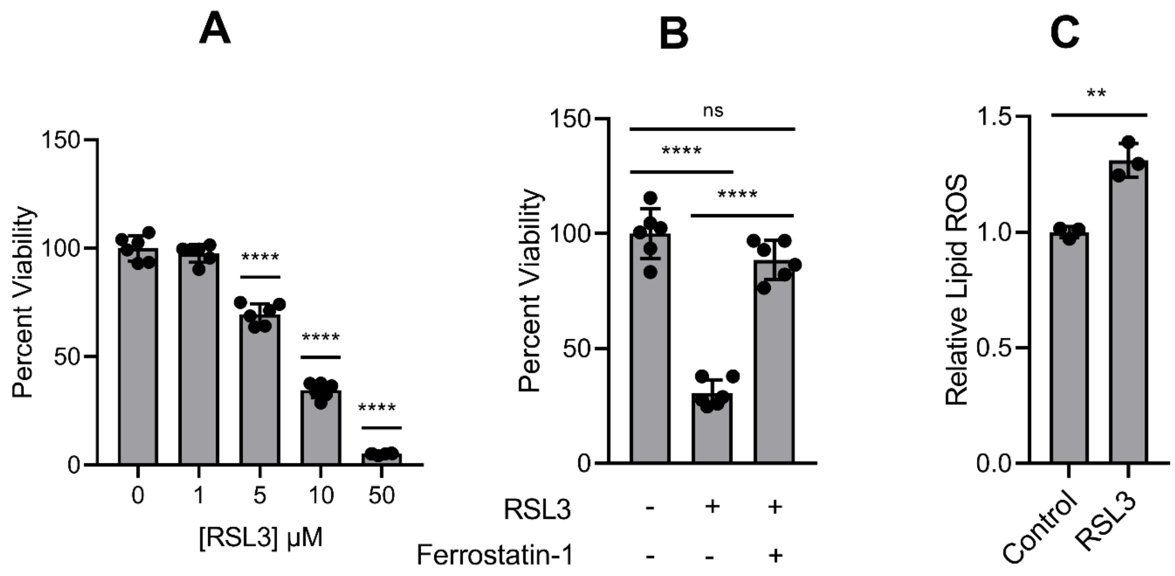

**Supplementary Figure 2.** Ferroptosis signaling pathway changes in RSL3-treated MC38 cell lines

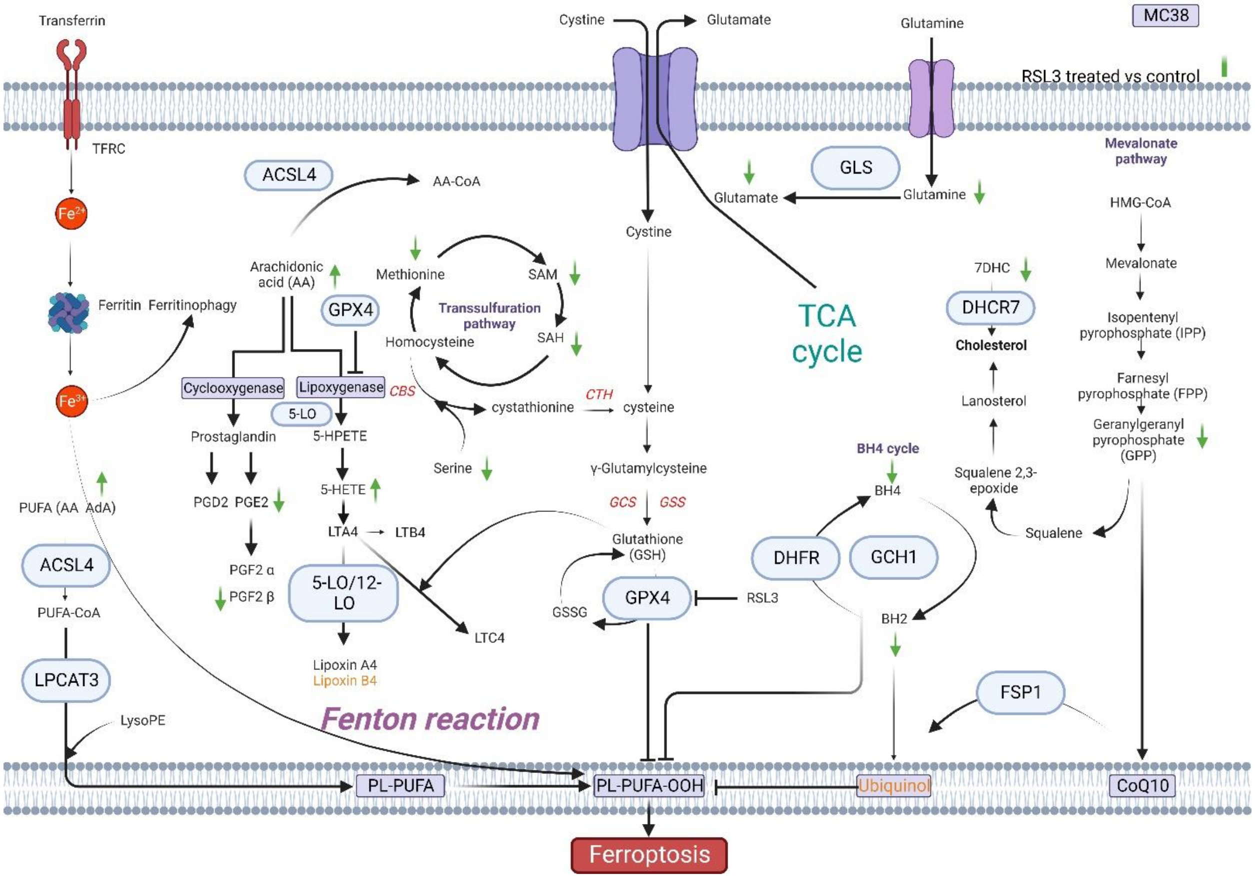


**Abbreviations:** ACSL4, acyl coenzyme A synthetase long-chain family member 4; BH2, dihydrobiopterin; BH4, tetrahydrobiopterin; CoA, coenzyme A; Fe, iron; FSP1, ferroptosis suppressor protein 1; GCH1, GTP Cyclohydrolase 1; GCS, glutamylcysteine synthetase; GLS, glutaminase; GPX4, glutathione peroxidase 4; GSH: reduced glutathione, GSSG: oxidized glutathione, GSS, glutathione synthetase; HMG-CoA, 3-hydroxy-3-methylglutaryl CoA; LPCAT3, lysophosphatidylcholine acyltransferase 3; NRF2, nuclear factor E2-related factor 2; PE, piperazine erastin; PL, phospholipid; PUFA, polyunsaturated fatty acid; TCA, tricarboxylic acid; LPCAT, lysophosphatidylcholine acyltransferase; PUFA, polyunsaturated fatty acids, MUFA, Monounsaturated fatty acids ; TFRC, Transferrin Receptor; PGD2, Prostaglandin D2; PGE2, Prostaglandin E2; PGF2α, Prostaglandin F2α; PGF2β, Prostaglandin FF2β; LTA4, Leukotriene A4; LTB4, Leukotriene B4; LTC4, Leukotriene C4; SAM, S-Adenosylmethionine; SAH, S-Adenosylhomocysteine; DHFR, Dihydrofolate reductase; DHCR7, 7-Dehydrocholesterol reductase; AA, Arachidonic acid; AdA: Adrenic acid

**Supplementary Figure 3.** Kaplan–Meier survival curves and *FTL* expression levels (low vs high) in tumors from CRC patients by sex and KRAS mutation type. Gene expression data obtained from NCBIs GEO GSE39582 dataset.

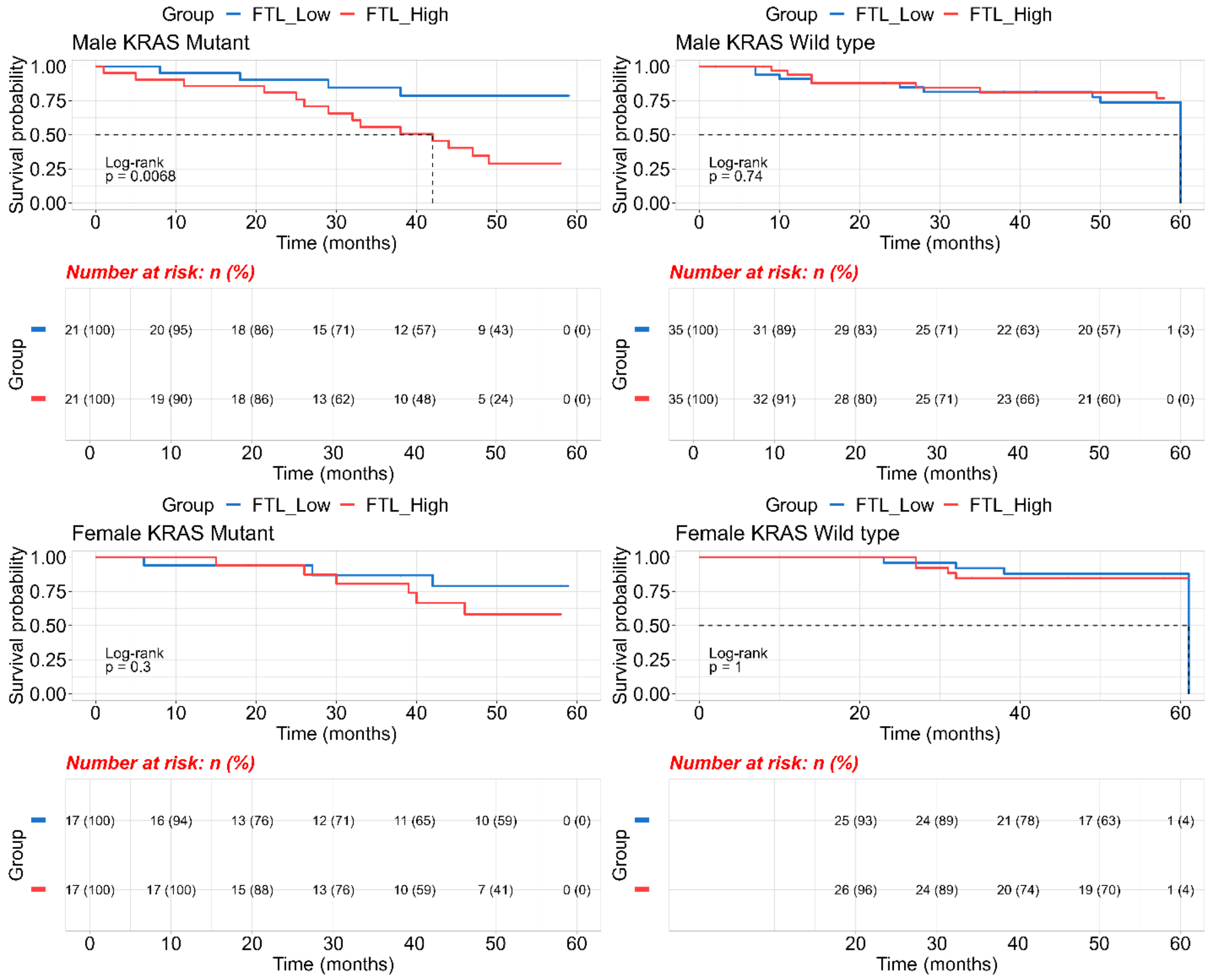

**Supplementary Figure 4.** Kaplan–Meier survival curve and *FTH1* expression levels (low vs high) in tumors from CRC patients by sex and KRAS mutation type. Gene expression data obtained from NCBIs GEO GSE39582 dataset.

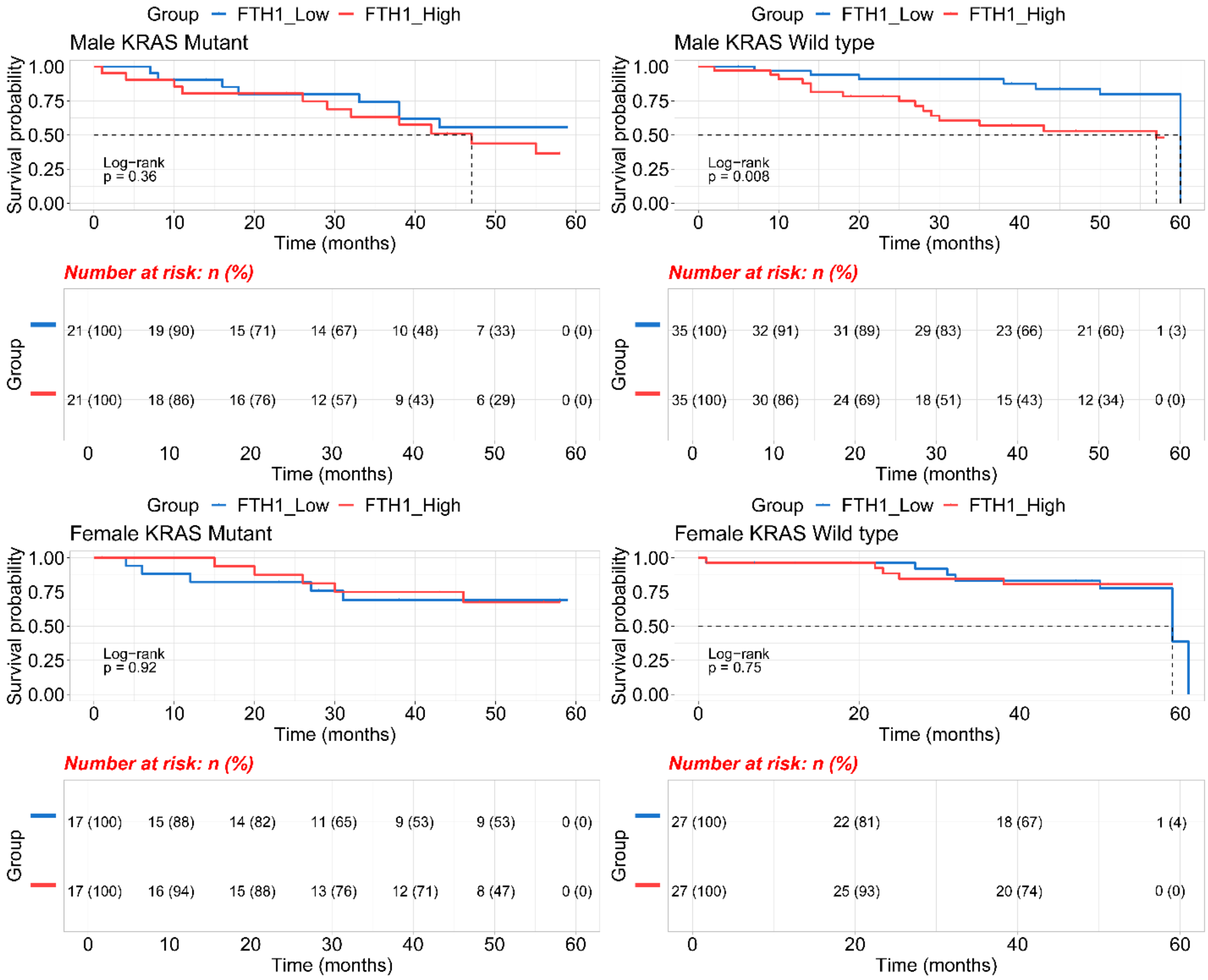

**Supplementary Figure 5.** Kaplan–Meier survival curve and *ASCL4* expression levels (low vs high) in tumors from CRC patients by sex and KRAS mutation type. Gene expression data obtained from NCBIs GEO GSE39582 dataset.

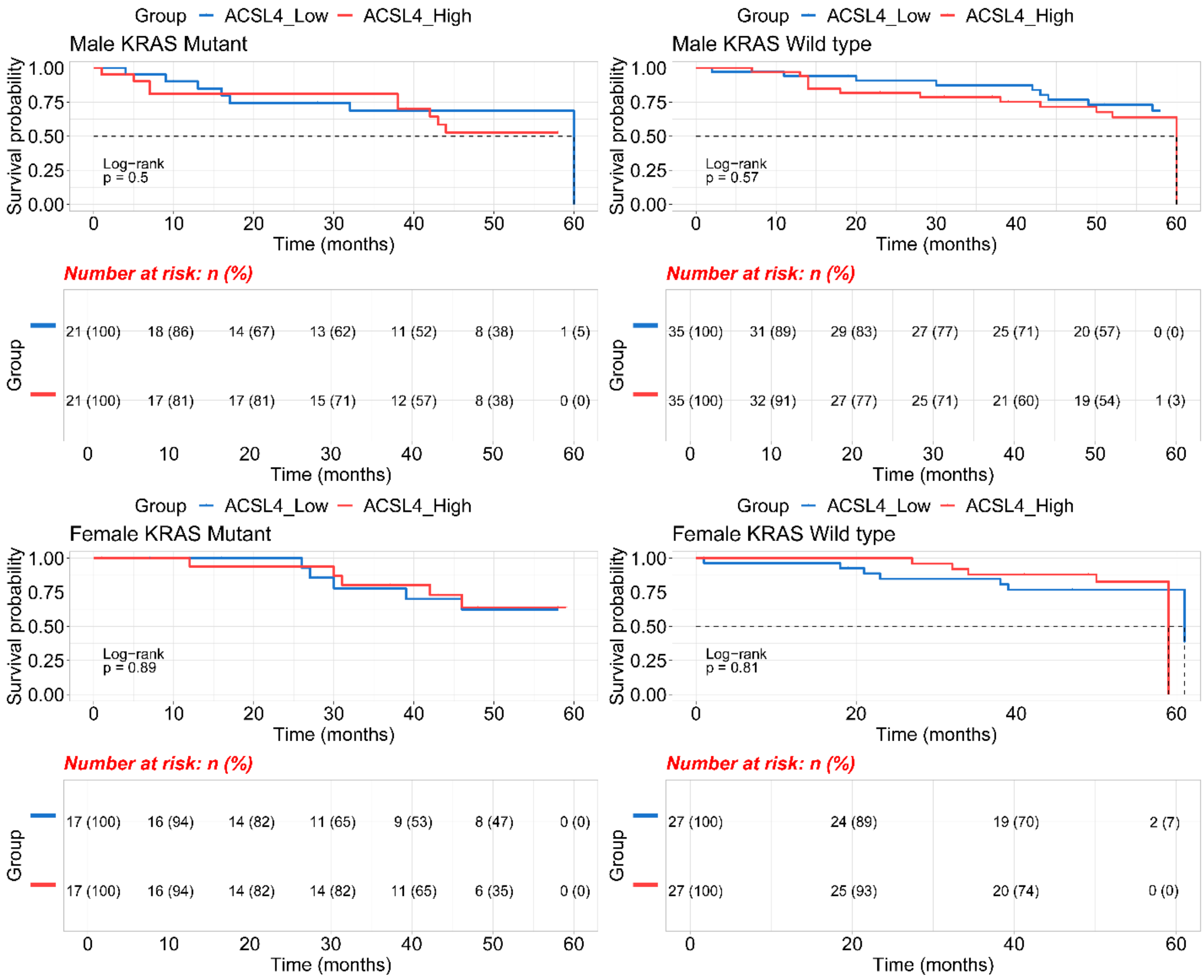

**Supplementary Figure 6**. Sex differences in genes associated with CRC prognosis. Hazard ratios (HRs) for 5-year overall survival (OS) by sex for (log2-transformed) gene expression, adjusted for anatomic location, chemotherapy history, clinical stage, and age (continuous), acquired by multivariate Cox PH regression. A gene with HR < 1 was associated with a protective effect on prognosis; gene with HR > 1 was associated with an adverse effect on prognosis. Genes with confidence intervals (CIs) marked with asterisks were significantly associated with the presented prognosis (Raw p values < 0.05). The x-axes are log-scaled. Significant sex interaction p values < 0.05.

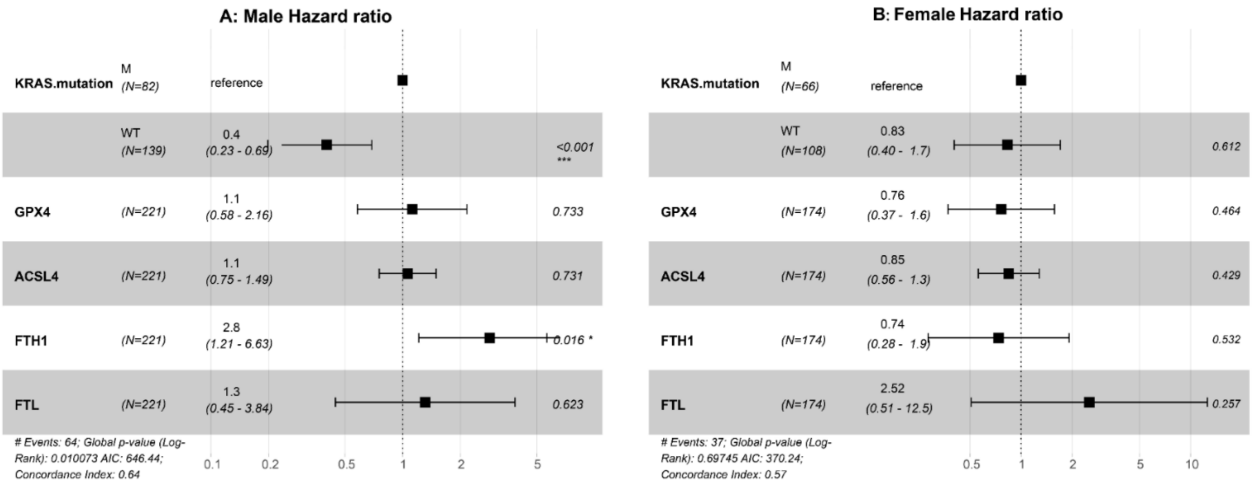
